# Physically Grounded Generative Modeling of All-Atom Biomolecular Dynamics

**DOI:** 10.64898/2026.02.15.705956

**Authors:** Bin Feng, Jiying Zhang, Xinni Zhang, He Cao, Ming Zhang, Patrick Barth, Zijing Liu, Yu Li

## Abstract

Predicting the kinetic pathways of biomolecular systems at all-atom resolution is crucial for understanding protein function and drug efficacy, yet this task is hindered by the immense computational cost of conventional molecular dynamics (MD) simulations. While deep learning has revolutionized static structure prediction and equilibrium ensemble sampling, simulating the kinetics of conformational transitions remains a critical challenge. We introduce BioKinema, a physically grounded generative model that predicts continuous-time, all-atom biomolecular trajectories at a fraction of the cost of traditional simulations. In particular, BioKinema utilizes a spatial-temporal diffusion architecture motivated by the exponential decay of correlations characteristic of Langevin dynamics, and is explicitly trained to generate trajectories with the correct kinetics and thermodynamics. It employs a hierarchical forecasting-and-interpolation strategy to overcome the error accumulation that often plagues long-horizon generation. Through extensive validation, we demonstrate that BioKinema generates physically stable and dynamically accurate trajectories suitable for rigorous downstream analysis. For protein systems, it reproduces the equilibrium thermodynamics of the conformational ensemble and the underlying kinetics. For protein-ligand complexes, it successfully elucidates mechanisms such as ligand-driven conformational changes and allosteric interactions. Furthermore, BioKinema leverages enhanced sampling data to predict rare kinetic events, emerging as a powerful tool for estimating ligand unbinding pathways. Collectively, these results establish BioKinema as a computationally efficient complement to MD simulations that bridges the gap between static structure and dynamic function, enabling high-throughput exploration of the kinetic landscape for structural biology and drug discovery.

## 1 Main

Biological function emerges not merely from static architecture but from conformational dynamics, where proteins must transition between distinct conformational states to drive biological processes^1^. Crucially, the kinetics of these transitions, which represent the continuous evolution between functional states, govern critical processes such as ligand binding^2^, signal transduction^3^, and enzymatic catalysis^4, 5^. Deciphering these mechanisms is therefore important for understanding the molecular basis of life. While recent computational breakthroughs, such as RoseTTAFold^6^ and AlphaFold^7, 8^, have revolutionized the prediction of static structures from sequence, knowledge of the ground state alone is often insufficient to capture the time-dependent behaviors, such as how ligands bind and unbind or allosteric signals propagate^9^. Consequently, bridging the gap between static structural models and the temporal evolution of dynamical molecular systems remains a central challenge.

In addressing these challenges, both experimental and computational approaches have been pursued. Experimental techniques such as stopped-flow spectroscopy^10^ or NMR^11^ can probe kinetic rates. However, they typically lack the spatial and temporal resolution to provide a complete, all-atom picture of the underlying dynamic pathways. Currently, Molecular Dynamics (MD) simulation is the primary computational method for exploring biomolecular kinetics at all-atom resolution^12, 13^. By numerically integrating equations of motion, MD can, in principle, reveal the complete time evolution of a molecular system. However, the practical utility of MD is severely hampered by its computational intensity^14^. Since femtosecond-scale time steps are required to resolve high-frequency atomic vibrations, the simulation of biologically relevant timescales can require days or months of computation^15^. While enhanced sampling methods like metadynamics^16, 17^ can accelerate the exploration of specific rare events, they often require predefined collective variables and remain computationally demanding for large systems^18^. These fundamental constraints have sustained a long-standing need for new approaches that can access long-timescale biomolecular kinetics at atomic detail without the prohibitive cost of brute-force simulation.

Generative machine learning offers a fundamentally different strategy. Already, structure prediction methods have demonstrated that static protein structures could be predicted from sequences with near-experimental accuracy^7, 8^, and generative models such as BioEmu^19^ subsequently extended this paradigm to thermodynamic equilibrium conformational ensembles^20–22^, with a primary focus on the backbone structures of single-chain proteins. Each successive advance has added a new dimension of physical completeness, progressing from ground-state snapshots to Boltzmann-weighted conformational distributions^23, 24^. The natural next frontier is the temporal dimension itself^25, 26^. This demands models capable of predicting not only which conformational states a biomolecule can access, but also the kinetic pathways and characteristic timescales governing transitions between them. Initial efforts toward time-aware generative models have demonstrated early promise. However, existing approaches are largely confined to single-protein systems^27, 28^ or rely on simplifications such as rigid-receptor approximations^29^, limiting their applicability for rational drug design such as allosteric drug targeting^9, 30^. The general challenge of generating physically plausible, all-atom kinetic trajectories for multi-component biomolecular systems remains open.

Here we introduce BioKinema, the first all-atom generative model designed to simulate long-timescale biomolecular dynamics with continuous time resolution. BioKinema is trained on approximately 54,600 *μ*s of aggregated MD trajectories. Leveraging the robust structural representations of Protenix^31^, the open-source reproduction of AlphaFold 3, BioKinema incorporates a novel Spatial-Temporal Diffusion Module with a physically motivated temporal attention mechanism. Its continuous-time attention bias *B*_*ij*_ = −*λ* |*t*_*i*_ − *t*_*j*_|^*β*^ is inspired by the Ornstein-Uhlenbeck process and enables the model to generalize across arbitrary time intervals^32^. To approximate the physical behavior of the biomolecular system, we explicitly train BioKinema to capture both the kinetics and thermodynamics properties of the system. Furthermore, to mitigate error accumulation in long-horizon generation, BioKinema adopts a unified “noise-as-masking” paradigm, enabling a hierarchical forecast-then-interpolate sampling strategy.

We validate the efficacy of BioKinema through extensive experiments spanning equilibrium thermodynamics, conformational kinetics, and rare unbinding events. Our results demonstrate that BioKinema consistently generates physically plausible structures and maintains trajectory stability up to microseconds, effectively circumventing the error accumulation issue of generative models. Building on this stability, we show BioKinema’s ability to reproduces the conformational kinetics and equilibrium thermodynamics of protein dynamics, recovering both the relative timescales of transitions between metastable states and their stationary populations. For protein-ligand systems, BioKinema accurately recovers intermolecular interactions and successfully resolves ligand-driven conformational changes and long-range allosteric interactions. Finally, we show that when fine-tuned on metadynamics data, BioKinema can predict rare kinetic events, such as ligand unbinding pathways, which remain computationally prohibitive for standard simulations. Collectively, these results establish BioKinema as a computationally efficient complement to conventional MD simulations, enabling rapid exploration of kinetic landscapes in structural biology and drug discovery.

## Results

### Overview of BioKinema

BioKinema is a generative model designed to simulate long-timescale, all-atom biomolecular dynamics with continuous time resolution. To address these existing challenges (**Figure 1a**), BioKinema enables fully flexible dynamics generation across diverse biomolecular systems including protein systems and protein-ligand complexes (**Figure 1b**). The model is trained on approximately 54,600 *μ*s of aggregated molecular dynamics trajectories, encompassing diverse biomolecular types and trajectory lengths ranging from nanoseconds to microseconds.

**Figure 1.**
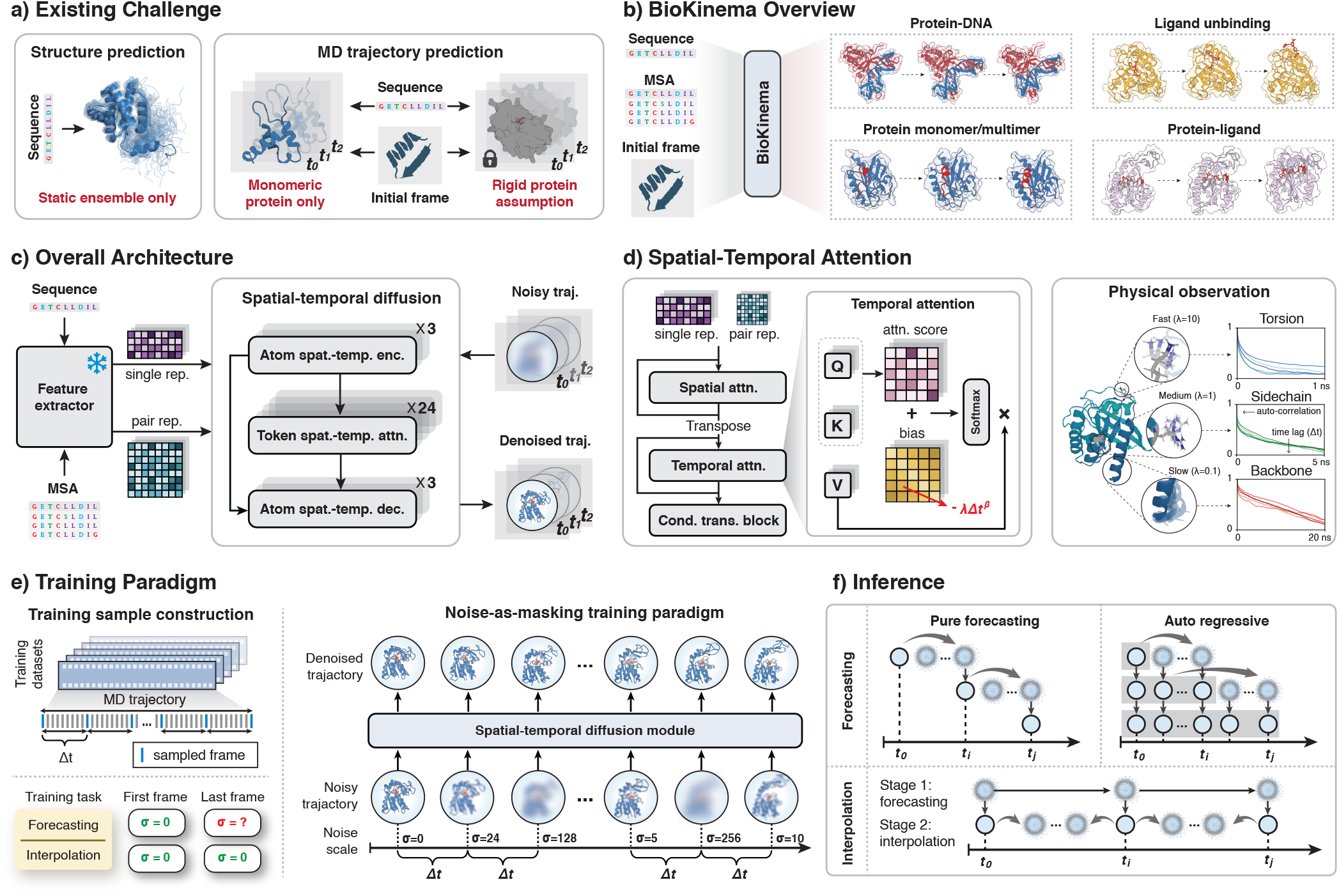
BioKinema: A physically grounded model for all-atom biomolecular dynamics generation. **(a)** Limitations of existing methods. Structure prediction models generate only static ensembles, while previous MD trajectory models are restricted to monomeric proteins or treat protein receptors as rigid bodies. **(b)** BioKinema enables fully flexible, all-atom dynamics generation for diverse biomolecular systems. **(c)** Overall model architecture of BioKinema. **(d)** Spatial-Temporal attention mechanism. A continuous-time attention bias encodes exponential decay of inter-frame correlations motivated by Langevin dynamics, and naturally captures dynamics on different time scales. **(e)** Unified “noise-as-masking” paradigm enabling a single model to perform both trajectory forecasting and interpolation. **(f)** Hierarchical sampling strategy combining coarse forecasting with fine-grained interpolation for long-horizon trajectory generation.

The overall architecture of BioKinema consists of two main components (**Figure 1c**): a frozen encoder comprising the Input Embedder and Pairformer modules from Protenix^31^ that generates single and pair representations, and a novel Spatial-Temporal Diffusion Module that generates continuous all-atom trajectories. The frozen encoder leverages rich structural and evolutionary priors from pre-trained weights while reducing training costs. Two key designs enable BioKinema to capture the physics of the underlying dynamics. First, we employ a physically motivated Spatial-Temporal Attention mechanism with a temporal bias motivated by Langevin dynamics (**Figure 1d**). In the overdamped limit, biomolecular coordinates follow an Ornstein-Uhlenbeck process with exponentially decaying temporal correlations, which we encode as a continuous-time bias term *B*_*ij*_ = −*λ*|*t*_*i*_ − *t*_*j*_|^*β*^ that yields stretched-exponential decay of the attention weights. Second, we train BioKinema with a physically grounded dynamics objective which supervises the transition rates between metastable states and their equilibrium populations, enabling BioKinema to capture the kinetic and thermodynamic properties of the biomolecular system (see **Supplements**).

To address the fundamental challenge of error accumulation in long-trajectory generation^33^, we make two complementary design choices. First, we introduce a noise-as-masking training paradigm (**Figure 1e**) which reframes diffusion noise levels as information visibility controls: clean frames (*σ* = 0) serve as fully visible conditioning inputs, while noised frames are prediction targets. By randomly designating certain frames as clean during training, a single unified model learns both forecasting and interpolation capabilities without requiring separate encoders or conditioning mechanisms. Second, this unified framework enables our hierarchical sampling strategy (**Figure 1f**), which decomposes long-trajectory generation into two stages. The first stage performs coarse-grained forecasting at large time intervals (e.g., 5–20 ns), establishing the global trajectory evolution. The second stage then fills in fine-grained details through interpolation, conditioning on the coarse trajectory frames as fixed anchors. This hierarchical approach enables BioKinema to efficiently generate microsecond-scale trajectories at arbitrary temporal resolutions while mitigating error accumulation problems.

### BioKinema Generates Physically Stable and Dynamically Accurate Trajectories

A foundational requirement for a generative MD model is the ability to produce trajectories that satisfy fundamental physical constraints. Consequently, we begin by evaluating whether BioKinema generates physically valid structures and reproduces the distributional properties observed in classical MD simulations. To rigorously assess the physical stability of generated trajectories for both protein and ligand components, we curated a challenging subset from the MISATO test set^14^, called MISATO-OOD, comprising 36 protein-ligand systems with a maximum sequence identity of 40% to the training set (**Supplementary Figures 1, 2**).

It is well known that error accumulation poses a significant challenge for the generation of long, physically stable trajectories^33^. To address this, we generated 1 *μ*s trajectories for each test system and systematically evaluated the physical integrity using three categories of metrics: (1) overall stability, evaluated by the Root Mean Square Deviation (RMSD) before and after energy minimization with the Amber99SB+GAFF2 force field; (2) ligand geometry, assessed by bond length and bond angle errors; and (3) protein quality, evaluated by MolProbity^34^ scores and the percentages of Ramachandran-favored backbone conformations and rotamer-favored side chain conformations. The results demonstrate that trajectories generated by BioKinema closely match MD simulations across all physical-fidelity metrics (**Figure 2a**). For overall stability, BioKinema-generated conformations exhibit a mean post-relaxation RMSD of 0.73 Å, close to the 0.69 Å in MD simulations. Regarding ligand geometry, BioKinema achieves a mean absolute error (MAE) of 0.046 Å for bond lengths and 0.056 rad for bond angles across all heavy-atom bonds, which closely aligns with the MD reference (0.042 Å and 0.048 rad). Furthermore, protein quality assessment shows a MolProbity score of 1.72 for BioKinema, comparable to the MD reference (1.43). Notably, when grouped by 0.1 *μ*s intervals, all metrics remained stable over the entire 1 *μ*s duration, with relative differences of at most 5.4% across subsequent time spans. These results confirm that BioKinema generates physically stable trajectories and effectively mitigates the error accumulation problem common in trajectory generation models, providing confidence for applying BioKinema to extended kinetic analyses.

**Figure 2.**
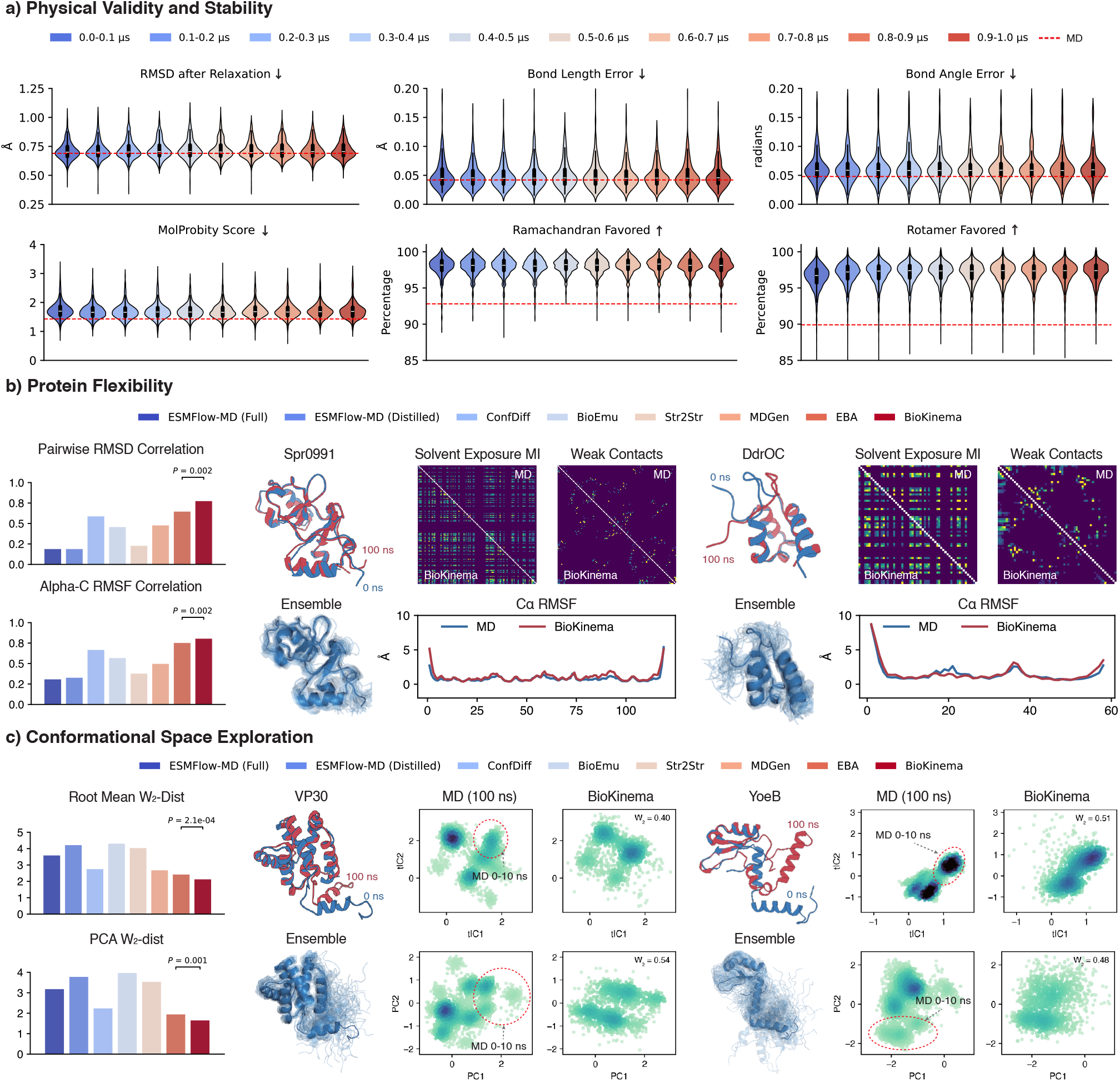
BioKinema Generates Physically Stable and Dynamically Accurate Trajectories. **(a) Physical validity and stability**. Violin plots of six stability metrics evaluated on 36 protein-ligand systems from the MISATO-OOD test set (max sequence identity to training ≤40%), each with 1 µs trajectories analyzed per 100 ns intervals. RMSD after energy minimization is computed with Amber99SB+GAFF2. (b) Protein flexibility. Left: Bar plots comparing BioKinema and competing methods on 82 single-chain proteins from the ATLAS test set in terms of pairwise RMSD and C*α* RMSF correlations. Right: Two cases (spr0991 and ddrOC) showing structural ensembles, solvent exposure mutual information maps, weak contact maps (contact probability (≤8 Å) ≤50%), and per-residue C*α* RMSF profiles comparing MD with BioKinema predictions. **(c) Conformational space exploration**. Left: Bar plot comparing BioKinema and competing methods on 82 single-chain proteins from the ATLAS test set in terms of Root Mean *W*_2_-Distance and PCA *W*_2_-Distance. Right: Two cases (VP30 and yoeB) showing conformations at 0 ns (input), 100 ns, and the complete generated ensemble, with tICA and PCA projections comparing 100 ns MD with BioKinema trajectories. Red dashed circles indicate MD 0–10 ns distribution. **(b**,**c)** All showing examples have a max sequence identity ≤40% to the training set.

Beyond physical stability, accurately capturing residue-level flexibility patterns is essential for understanding protein function, as local dynamics govern critical processes such as enzyme catalysis, ligand binding, and allosteric regulation^2, 18, 35^. To evaluate this capability, we benchmarked BioKinema against seven competing methods on the ATLAS^36^ test set (82 single-chain proteins), using pairwise RMSD and C*α* RMSF correlations to assess per-target and per-residue flexibility, respectively. For each test system, we generated 10 independent 100 ns trajectories from each MD initial frame. BioKinema significantly outperforms all competing methods (**Figure 2b**), achieving a pairwise RMSD correlation of approximately 0.80, compared to 0.65 for the second-best method, EBA^37^. Similarly, for C*α* RMSF correlation, BioKinema reaches 0.82, surpassing the 0.62 for EBA. These results indicate that BioKinema not only captures global conformational diversity but also faithfully predicts local flexibility profiles. In addition, our ablation study confirms the effectiveness of our proposed hierarchical sampling strategy and the positive contribution of the MSA module (**Supplementary Figure 3**). The performance of BioKinema extends to the ATLAS-OOD test set, which comprises 59 functionally distinct proteins with ≤ 40% sequence identity to the training set. As shown in **Figure 2b** and **Supplementary Figures 4–6**, the C*α* RMSF profiles of BioKinema closely match the MD ground truth, accurately reproducing high flexibility peaks. Furthermore, an analysis of correlated motions via solvent exposure mutual information (MI) maps reveals a correlation of 0.59 between MD and BioKinema, indicating that BioKinema correctly captures the dynamic coupling of solvent accessibility, a feature essential for domain motions and allosteric signaling. Besides, the weak contact maps further demonstrate that BioKinema accurately reproduces the distribution of short-lived contacts that drive critical conformational transitions, such as van der Waals interactions and transient hydrogen bonds, with a correlation of 0.55 between MD and BioKinema. Collectively, these results indicate that the generated ensembles are statistically consistent with physical MD trajectories, rather than merely matching average structures.

Finally, while accurate flexibility patterns indicate local correctness, a comprehensive MD emulator must also explore the global conformational landscape faithfully, which is crucial for discovering rare but functionally relevant states. To assess this, we evaluated the distribution similarity between generated and ground-truth ensembles across 82 proteins in the ATLAS test set using the Wasserstein-2 (*W*_2_) distance. As illustrated in **Figure 2c**, BioKinema achieves a Root Mean *W*_2_-Distance of ∼ 2.3 (in raw coordinate space) and a PCA *W*_2_-Distance of ∼ 1.8 (in the principal component projection space), representing improvements of ∼ 20% and ∼ 28% over the second-best method, respectively. These results demonstrate that BioKinema most faithfully reproduces the global conformational distribution of MD simulations. To further visualize this capability, we examined two highly flexible proteins with low sequence identity (≤ 40%) to the training set: VP30, an Ebola virus transcription activator, and YoeB, a bacterial toxin. In the tICA (lag time = 1 ns) and PCA projections (**Figure 2c**), the BioKinema-generated ensembles overlap well with the conformational space of the full 100 ns MD trajectory, while the initial 0–10 ns MD simulations (red dashed circles) remain confined to a smaller space. Importantly, BioKinema generates these 100 ns MD trajectories in less than one minute on a single GPU, showing a substantial efficiency advantage over the approximately 22 hours required for equivalent conventional MD simulations^38^.

### BioKinema reproduces conformational kinetics and equilibrium thermodynamics

A defining strength of molecular dynamics is that it provides two complementary properties that are inaccessible to static structures: the equilibrium *thermodynamics* of the conformational ensemble, and the conformational *kinetics* governing transitions between metastable states. These two aspects are closely related but probe distinct properties of a system. Thermodynamic observables assess whether the correct equilibrium ensemble is recovered, whereas kinetic observables further probe whether the temporal evolution within that ensemble, the rates and pathways of interconversion between states, is faithfully reproduced.^39^ We therefore asked whether BioKinema recovers not only the plausible structures, but also their underlying thermodynamic and kinetic behavior.

To answer this question, we constructed the CATH2-OOD benchmark of 20 held-out protein domains from the CATH2 test set (sequence identity ≤ 40% to the training set), each with 32–41 independent 1 *μ*s MD reference trajectories at 10 ns resolution. For every MD trajectory, we generated 5 independent BioKinema trajectories from its initial frame, also 1 *μ*s at 10 ns resolution, yielding approximately 3,800 generated trajectories in total. Both MD and BioKinema ensembles were projected onto the same time-lagged independent component (TICA) space^40, 41^ and discretized using the same *K* = 10 Markov state model (MSM) constructed from MD^39^, enabling direct comparison of kinetic and thermodynamic observables.

We first evaluated whether BioKinema reproduces the conformational kinetics of MD. We quantified transition dynamics using the pairwise mean first-passage times (MFPTs) between MSM states. Across all 20 systems, BioKinema accurately recovers which transitions are fast and which are slow (**Figure 3a**, left), with a mean log-space Pearson correlation of 0.77 between the BioKinema and MD MFPTs across transitions spanning three orders of magnitude in timescale (10^1^–10^5^ ns). The agreement extends to slow collective dynamics: the mean absolute error (MAE) between BioKinema and MD TICA autocorrelation-function (ACF) is 0.075, and the mean relaxation time *τ*_1/e_ of the slowest component is 530 ns for BioKinema versus 539 ns for MD. These results indicate that BioKinema captures not only individual conformations but also the long-timescale dynamical processes that govern conformational switching, demonstrating that BioKinema reproduces the kinetics of the underlying dynamics. Representative systems further illustrate this behavior (**Figure 3a**, right). For both FADD and carB, BioKinema reproduces the major conformational basins sampled by MD, closely matches the ordering of pairwise MFPTs (log-space Pearson correlations of 0.72 and 0.80, respectively), and accurately recovers the relaxation dynamics of the slowest collective coordinates (e.g., 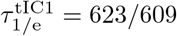 ns for BioKinema/MD on FADD).

**Figure 3.**
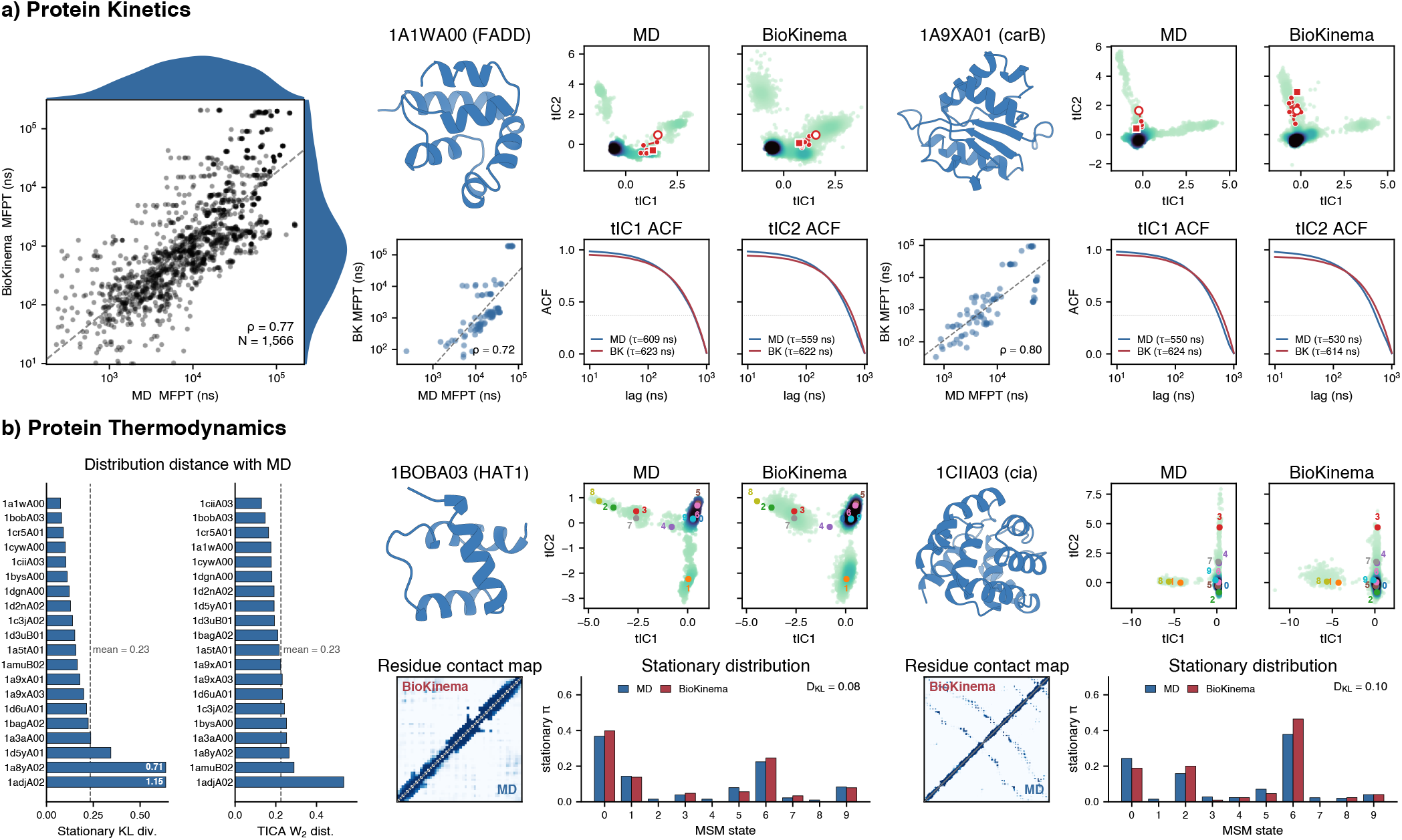
BioKinema reproduces conformational kinetics and equilibrium thermodynamics. **(a) Protein kinetics**. Left: Joint plot showing mean first-passage times (MFPT) between *K*=10 Markov state model (MSM) states of MD and BioKinema on the 20 CATH2-OOD test systems (*N* = 1,566 MSM state pairs). Right: Two case studies (FADD and carB), each showing the protein structure, density-coloured time-lagged independent component (TICA) ensembles with a representative trajectory (red), scatter plot showing per-protein MFPT, and line plot showing TICA-1 / TICA-2 autocorrelation functions (ACF) for MD (blue) and BioKinema (BK, red) with the relaxation time *τ*_1/e_ annotated. **(b) Protein thermodynamics**. Left: Per-system KL divergence (*D*_KL_) between MD and BioKinema MSM stationary distributions, and root-mean Wasserstein-2 (*W*_2_) distance over the five slowest TICA dimensions, sorted in ascending order. Right: Two case studies (HAT1 and cia), each showing the protein structure, density-coloured TICA ensembles with the MSM cluster centres, the C*α* contact-probability map with 8 Å contact cutoff (mean absolute errors 0.012 and 0.003 for HAT1 and cia), and bar plot showing the MSM stationary distribution over *K*=10 states with *D*_KL_ annotated. **(a, b)** BioKinema generates 5 trajectories (1 *µ*s with 10 ns time interval) from each of 32–41 MD initial frames per system.

Having established the kinetic fidelity of BioKinema, we next asked whether these accurate dynamics also give rise to the correct equilibrium thermodynamics. We first compared the MSM stationary distributions of BioKinema and MD directly, quantifying their agreement by the Kullback–Leibler divergence (*D*_KL_). BioKinema matches the MD equilibrium closely across the test set (**Figure 3b**, left): 18 of 20 systems achieve *D*_KL_ ≤ 0.4, with a mean of *D*_KL_ = 0.23 (**Supplementary Table 3**; per-system results in **Supplementary Figures 9–11**), indicating that the generated dynamics recover the correct equilibrium populations of the metastable states. Because the stationary distribution is defined over coarse-grained metastable states, we next compared the continuous equilibrium ensembles in the space of five slowest TICA dimensions using the root-mean Wasserstein-2 (*W*_2_) distance, obtaining a mean *W*_2_ = 0.23 across all systems. Finally, we assessed agreement directly in Cartesian space by comparing per-residue C*α* contact-probability maps: the mean absolute difference between BioKinema and MD is only 0.006 across all 20 systems. These three complementary analyses consistently demonstrate that BioKinema recovers the equilibrium conformational ensemble across discrete state populations, continuous slow collective coordinates, and all-atom structural observables. Two representative systems, HAT1 and cia, illustrate this agreement across all three measures (**Figure 3b**, right). An ablation study further confirms the contribution of the physically grounded dynamics loss, whose removal degrades every kinetic and thermodynamic metric across the test set (**Supplementary Figure 12**). Collectively, these findings establish that BioKinema faithfully reproduces both equilibrium thermodynamics and conformational kinetics.

### BioKinema captures protein-ligand interactions

Protein-ligand interactions are fundamental to nearly all biological processes and form the mechanistic basis for the majority of therapeutic interventions^42–44^. However, modeling these interactions remains a significant challenge for deep learning approaches due to the highly coupled dynamics between the ligand and the protein pocket. An effective model must not only capture the individual motions of the protein and ligand, but also reproduce their mutual interactions and the conformational changes induced by ligand binding. In this section, we evaluate the capacity of BioKinema to capture these complex dependencies and accurately reproduce the statistical ensembles of protein-ligand interactions.

To quantitatively assess interaction fidelity, we utilized the MISATO^14^ test set, comprising 1,031 diverse targets after filtering (see Methods). We employed Interaction Map Similarity as the primary metric, defined as the similarity between the protein-ligand contact probability maps generated by BioKinema and those derived from ground-truth MD simulations. To establish a baseline for intrinsic MD simulation variability, we also computed the similarity between two independent MD replicates (MD1 vs. MD2, 4 ns each). The results demonstrate that BioKinema achieves high interaction fidelity, with a mean Interaction Map Similarity of 0.88, compared to 0.91 between independent MD runs. As illustrated in the joint distribution plot (**Figure 4a**, left), the similarity between BioKinema and MD (y-axis) exhibits a strong positive correlation with the internal consistency of MD simulations (x-axis), indicating that BioKinema captures the interaction landscape with an accuracy comparable to classical MD.

**Figure 4.**
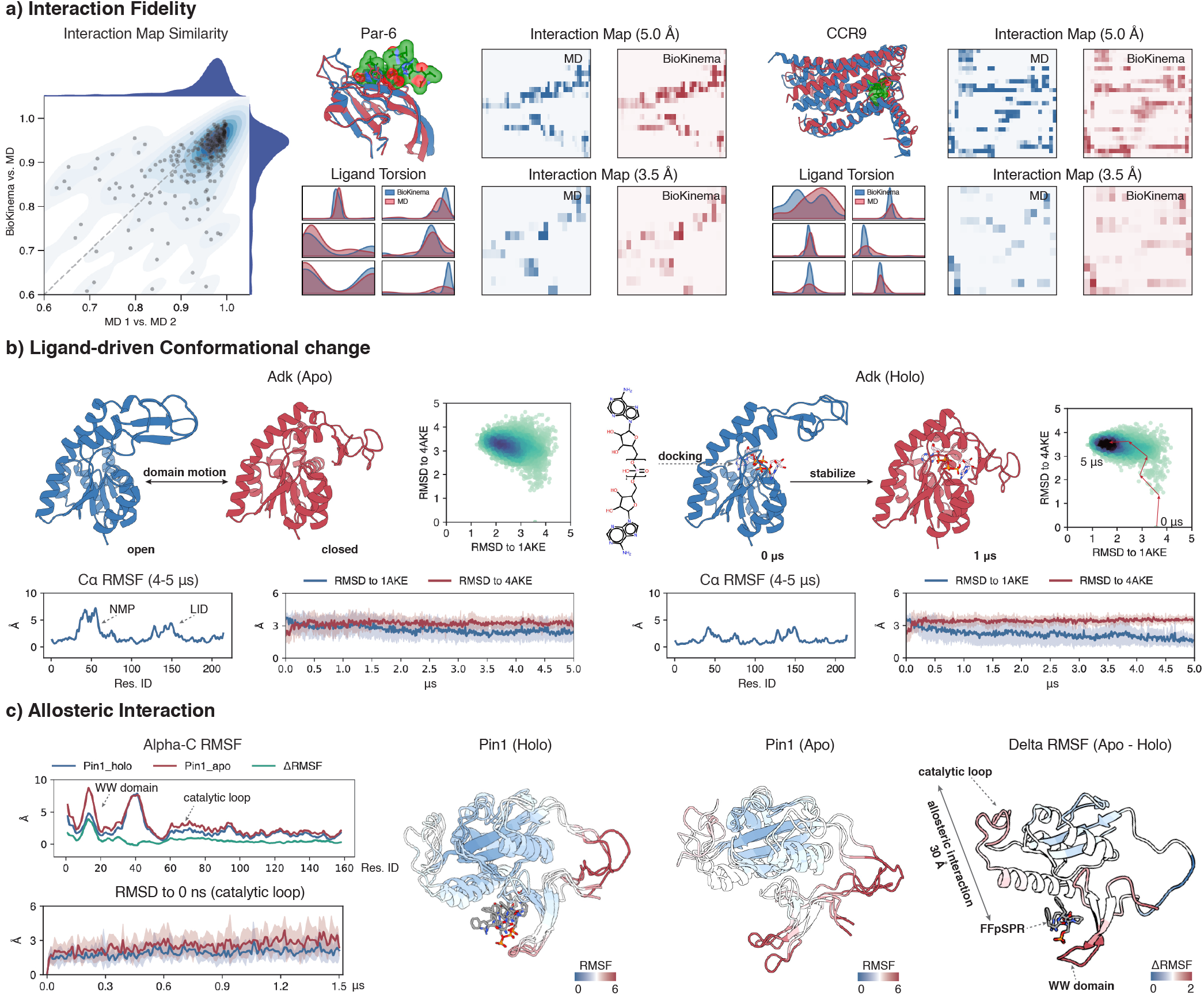
BioKinema captures protein-ligand interactions. **(a) Interaction fidelity.** Left: Joint plot comparing interaction map similarity between BioKinema vs. MD (8 ns) and MD1 (4 ns) vs. MD2 (4 ns) on the MISATO test set (1,031 targets). For each system, one 100 ns trajectory was generated to compute interaction maps. Right: Two case studies (Par-6 and CCR9) showing input structures (0 ns), generated conformations (100 ns), ligand torsion distributions, and interaction maps. **(b) Ligand-driven Conformational change**. Left: Adk (Apo) showing domain motion between open and closed states. Right: Adk (Holo) showing ligand-induced conformational change, the input structure is prepared using FitDock. Bottom: Line plots showing C*α* RMSF profiles and RMSD to initial structure. **(c) Allosteric interaction**. Left: Lines plot showing C*α* RMSF profiles and catalytic loop RMSD to the initial structure. Middle: Structural snapshots colored by RMSF for holo and apo states. Right: Structure colored by ΔRMSF (Apo − Holo). **(b, c)** All trajectories are generated 10 times independently. Protein conformations are initialized from 4AKE (Adk) and 3TDB (Pin1) during simulation. Shaded regions in the RMSD to initial structure plot indicate min-max range across 10 independent trajectories.

We further contextualize this performance through two representative case studies from the MISATO-OOD test set: the Par-6 PDZ domain, a cell polarity regulator involving surface peptide recognition, and the CCR9 receptor, a GPCR where the antagonist binds within a deep transmembrane pocket. For both systems, BioKinema successfully reproduced the interaction landscapes (**Figure 4a**, middle and right). Specifically, the interaction maps at both 5.0 Å (all contacts) and 3.5 Å (strong interactions) cutoffs show that BioKinema (red) accurately replicates the contact patterns observed in MD (blue). Furthermore, the ligand torsion angle distributions generated by BioKinema show substantial overlap with the MD-sampled ensembles, demonstrating that the model captures the conformational flexibility of the ligand within diverse binding site architectures, ranging from solvent-exposed surfaces to buried binding pockets (see **Supplementary Figures 7,8**).

A critical test for any generative dynamics model is the ability to simulate large-scale conformational changes driven by ligand binding. We investigated Adenylate Kinase (Adk), a classic model system that undergoes a transition between an “open” state (allowing substrate entry) and a “closed” state (catalytically active) via the movement of its LID and NMP domains^45^. We first simulated Adk in its ligand-free (Apo) state, initiating the trajectory from the open conformation (PDB: 4AKE). Consistent with NMR and computational studies for the apo state^46^, BioKinema’s simulation exhibited significant intrinsic flexibility (**Figure 4b**, left), with the Root Mean Square Fluctuation (RMSF) analysis showing high mobility in the LID and NMP domains. The RMSD distributions relative to both the open state (4AKE) and the closed state (1AKE) indicate that the protein spontaneously samples both conformations, transitioning between open and closed states in the absence of ligand.

In contrast, when the inhibitor AP5 was docked into the open structure (Holo state), BioKinema generated a markedly different trajectory that captured the conformational transition (**Figure 4b**, right). A rapid, ligand-driven transition was observed. The RMSD to the closed state (1AKE) decreased sharply within the first 2 *μ*s, while the RMSD to the open state (4AKE) increased. Following this transition, the RMSF profile showed a marked reduction in the fluctuations compared to the Apo state, indicating that the ligand effectively shifts the conformational equilibrium toward the closed state, consistent with NMR studies^46^. By successfully capturing the transition from a flexible open ensemble to a ligand-stabilized closed state, BioKinema demonstrates the capability to predict functional domain motions characteristic of induced-fit binding.

Finally, accurately modeling long-range allosteric regulation poses a fundamental challenge in computational biology^47–49^, as it requires the model to capture correlations between atoms separated by vast distances, rather than merely local physical interactions. To evaluate BioKinema’s ability to learn these long-range dependencies, we examined Pin1 (PDB: 3TDB), a peptidyl-prolyl isomerase where ligand binding at the interface between the WW domain and PPIase domain allosterically regulates the distal catalytic loop, located approximately 30 Å away^50^. In the Apo state, the catalytic loop of Pin1 is known to be semi-disordered^51^. BioKinema correctly reproduced this behavior, showing high C*α* RMSF values in the WW domain and the catalytic loop region. The RMSD of the catalytic loop relative to the initial structure increased rapidly during the simulation, reflecting the inherent flexibility of this region in the absence of ligand (**Figure 4c**, left).

Conversely, in the Holo state (bound to FFpSPR), BioKinema predicted a significant stabilization of the distal catalytic loop. The ΔRMSF analysis (Apo − Holo) reveals a clear allosteric pathway: ligand binding at the N-terminal WW domain transmits a stabilizing effect through the interdomain interface, ultimately reducing fluctuations in the C-terminal catalytic loop 30 Å away (**Figure 4c**, right). The RMSD of the catalytic loop in the Holo state remained low and stable throughout the simulation, aligning with the previous MD evidence^51^. This successful reconstruction of the Pin1 allosteric mechanism suggests that BioKinema captures global cooperative motions and long-range structural correlations, which are important considerations for understanding and designing allosteric modulators.

### BioKinema generates accurate ligand unbinding pathways

Ligand unbinding kinetics determines drug residence time, a key pharmacokinetic parameter. However, these dissociation events typically occur on timescales ranging from milliseconds to hours, computationally nearly inaccessible for conventional MD^17^. This timescale gap implies that models trained solely on equilibrium MD data are incapable of producing the rare, non-equilibrium unbinding pathways. To bridge this gap, we fine-tuned BioKinema on DD-13M^17^, a large-scale metadynamics database of ligand unbinding dynamics, which contains over 26,000 dissociation trajectories across 565 complexes. During training, we applied a causal mask due to the history-dependent nature of metadynamics sampling (see **Supplements**).

For evaluation, we curated a challenging test set of 31 protein-ligand systems with low sequence identity (≤ 40%) to the training set. We generated 20 unbinding trajectories for every test system using an auto-regressive denoising strategy, a method that incrementally incorporates generated frames into the context window (see **Methods**). To verify that the generated trajectories are physically plausible, we first assessed their structural stability, focusing on ligand stability metrics including bond length error, bond angle error, and protein–ligand clashes. As shown in **Figure 5a** and **Supplementary Figure 13**, BioKinema with auto-regressive generation closely matches metadynamics references across all evaluation metrics. For ligand geometry, BioKinema achieves a ligand bond error of 0.030 Å and a ligand angle error of 0.054 rad, comparable to the metadynamics references of 0.025 Å and 0.045 rad, respectively. Furthermore, when we grouped the test results by 6 ps intervals (60 frames), we observed that errors accumulate significantly over time due to the substantial conformational changes during ligand dissociation. Comparing the first and last time intervals, the model fine-tuned from BioKinema shows 51% and 55% increases in ligand bond error and ligand angle error, respectively, whereas the model trained from scratch exhibits 101% and 90% increases. This demonstrates that pre-training on equilibrium MD data provides geometric priors that mitigate error accumulation during extended trajectory generation.

**Figure 5.**
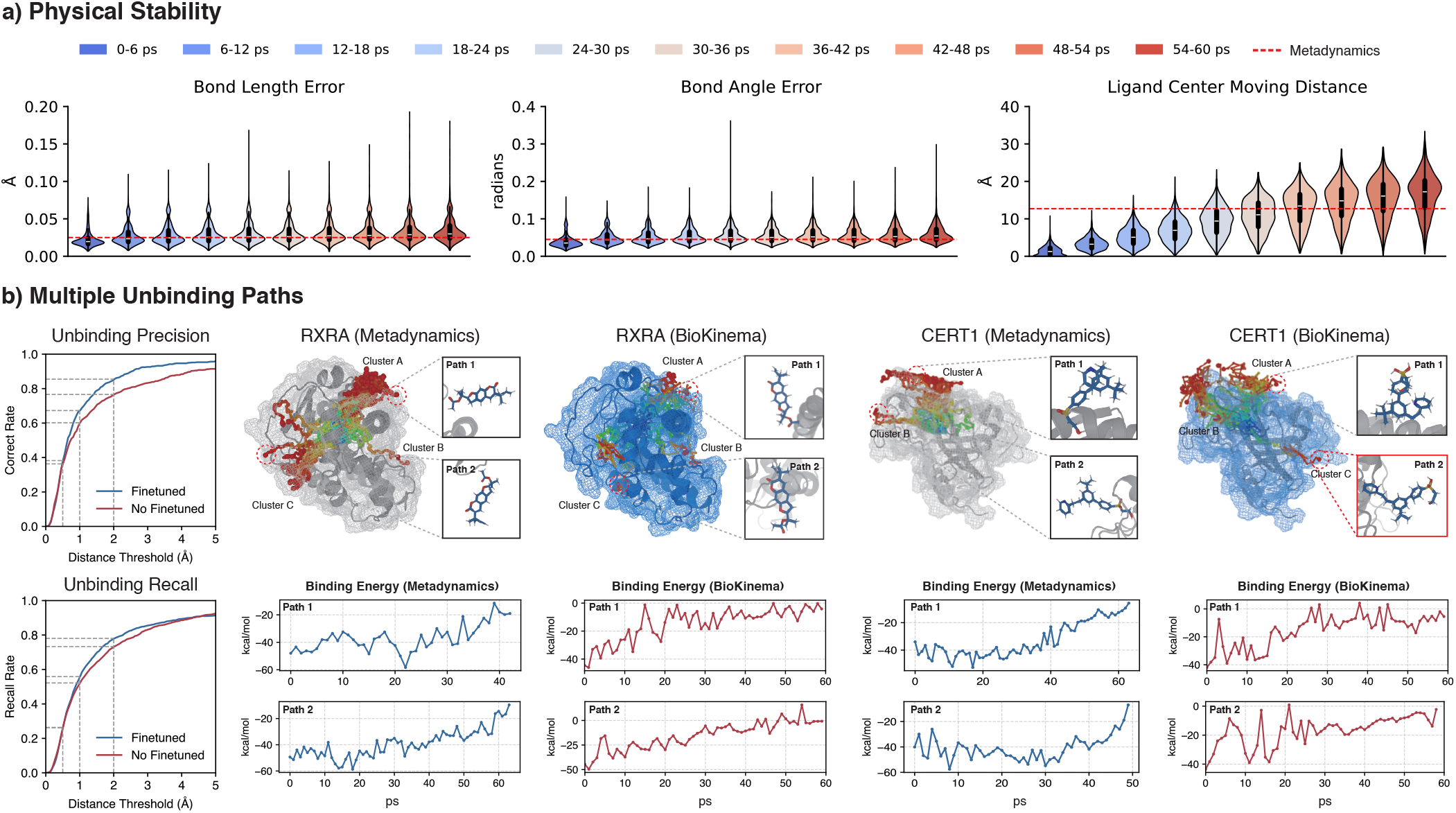
BioKinema generates accurate ligand unbinding pathways. **(a) Physical stability over extended simulations**. Violin plots showing bond length error, bond angle error, and ligand center moving distance across 60 ps trajectories, evaluated on the DD-13M test set (sequence identity *≤*40% to training set). **(b) Multiple unbinding pathways**. Left: Precision (correct rate) and recall curves as a function of endpoint distance threshold comparing model finetuned from BioKinema (blue) and non-finetuned (red), evaluated on the DD-13M test set. Right: Two case studies for RXRA (retinoic acid receptor) and CERT1 (ceramide transfer protein). Each unbinding path is colored from blue (bound state) to red (unbound state). A novel pathway discovered by BioKinema that was not observed in metadynamics is highlighted in red. Bottom: Line plots showing binding energy of representative trajectories calculated using GBSA. **(a, b)** All trajectories are generated 20 times independently using an auto-regressive method with 10 ps time step for each iteration.

Having established structural stability, we next evaluated the pathway accuracy of the generated unbinding trajectories. As a history-dependent bias potential artificially accelerates metadynamics, the resulting trajectories do not reflect physical residence times. Our evaluation therefore focuses on structural pathway accuracy. Analysis of the ligand’s geometric center reveals a progressive displacement from the binding pocket. Starting from zero, the displacement reached 16.63 Å in the 54–60 ps segment, confirming that the ligand moves away from the binding pocket. To quantify the pathway fidelity, we then computed precision and recall by measuring whether the generated unbinding trajectories end within a distance threshold of any metadynamics-discovered exit point (**Figure 5b** and **Supplementary Figure 14**). At a distance threshold of 2 Å, 87.4% of generated trajectories match at least one metadynamics-discovered pathway (precision), indicating high endpoint accuracy. Simultaneously, 77.7% of metadynamics trajectories are matched by at least one generated trajectory (recall), demonstrating broad coverage of the known dissociation routes.

When ligands occupy buried binding sites within structurally complex proteins, multiple unbinding pathways may exist. We present two case studies from our test set featuring multiple exit routes (**Figure 5b**, right). For the cases of RXRA (retinoic acid receptor alpha) and CERT1 (ceramide transfer protein), BioKinema successfully reproduced all pathway clusters identified by metadynamics, achieving an unbinding precision of 0.90 at a distance threshold of 2 Å. Furthermore, in the case of CERT1, the model identified a potential novel unbinding pathway not observed in the metadynamics simulations. To validate the energetic plausibility of the generated trajectories, we calculated the binding energy of each conformation using GBSA (Generalized Born and Surface Area solvation)^52^ (**Figure 5b**, bottom). The binding energy progressively increases along the trajectories and approaches zero as the ligand reaches the protein surface, consistent with the expected energetic behavior of dissociation. The novel pathway identified by BioKinema also exhibits binding energy profiles comparable to those of metadynamics-discovered pathways, supporting its physical plausibility. Collectively, these findings confirm that BioKinema can generate physically plausible unbinding trajectories that recapitulate known dissociation pathways and may identify additional routes not captured by metadynamics sampling.

## Discussion

In this work, we presented BioKinema, a generative foundation model capable of simulating all-atom biomolecular dynamics with continuous time resolution. Trained on a large-scale dataset spanning diverse biomolecular systems, BioKinema effectively captures both equilibrium fluctuations and non-equilibrium kinetic transitions across protein monomers and protein-ligand complexes. Our results demonstrate that a single, unified architecture can learn transferable representations of molecular dynamics, enabling accurate simulation of complex mechanistic phenomena, ranging from ligand-driven conformational changes and allosteric interactions to the rare kinetics of ligand unbinding.

There are key distinctions between BioKinema and preceding generative approaches. While time-agnostic models, such as BioEmu^19^, DiG^53^, and AlphaFlow^20^, successfully predict thermodynamic ensembles, they inherently discard the temporal dimension, failing to recover the transition pathways that define biological function. Conversely, current approaches that retain temporal order often impose constraints, such as the rigid-receptor assumption utilized in NeuralMD^29^. BioKinema addresses these limitations by modeling the protein and ligand as a coupled, fully flexible system. By explicitly approximating the temporal evolution of coordinates rather than generating independent snapshots, the model recovers kinetic properties essential for mechanistic analysis and effectively bridges the gap between static structural prediction and dynamic underpinnings of protein function.

Despite these promising results, several limitations remain. First, the simulation of events on extremely long timescales (on the order of milliseconds) remains computationally prohibitive^54^. While our hierarchical sampling effectively controls error accumulation at the microsecond scale, biological processes occurring at millisecond timescales often involve massive conformational rearrangements, such as global unfolding or large-scale domain swapping, where the model’s behavior has yet to be fully validated. Moreover, even with the major advances afforded by specialized hardware such as Anton^15^, millisecond-timescale trajectories remain too scarce to constitute sufficient training data in this dynamic regime. Second, unlike MD, BioKinema learns an empirical distribution from data rather than explicitly solving an energy function. Consequently, thermodynamic parameters such as temperature or chemical conditions such as pH and ionic strength are implicitly encoded within the model weights rather than being standard tunable variables as in classical MD^55^. Finally, BioKinema can access alternative conformations only through kinetic transitions between metastable states, and it learns these transitions directly from MD trajectories. Because MD itself faces a fundamental difficulty in crossing high free-energy barriers within a single trajectory, BioKinema inherits this same limitation; consequently, sampling a biomolecule’s full Boltzmann equilibrium distribution from any one trajectory remains difficult. We nonetheless offer an effective alternative: given a diverse set of starting conformations, BioKinema samples multiple independent replica trajectories that, when pooled via TICA+MSM analysis, accurately recover both the equilibrium thermodynamics and the kinetics (**Figure 3b**).

In conclusion, while BioKinema does not yet fully replace classical MD for scenarios requiring precise thermodynamic control, it represents a fundamental step toward efficient, generalizable exploration of the kinetic landscape of biomolecular motions. As high-quality trajectory data accumulates, we anticipate that foundation models for molecular dynamics will become increasingly useful tools for understanding the dynamic basis of biological function^56^.

## Methods

### Preliminaries

#### Notations

A biomolecular trajectory containing *T* frames of coordinates is denoted as **X**_*T*_ = {**x**_0_, **x**_1_, · · · **x**_*T* −1_} ∈ ℝ^*T* ×*N* ×3^, where **x**_*t*_ ∈ ℝ^*N* ×3^ represents the coordinates at time-step *t*, and *N* is the number of atoms in the biomolecular system. The trajectory prediction task aims to generate the sequence of conformations **x**_1_, …, **x**_*T* −1_ given an initial structure **x**_0_.

#### Diffusion models

Diffusion models are a class of generative models that learn to reverse a gradual noising process. In a Euclidean space, the forward process progressively corrupts clean data **x** ∼ *p*_data_ by adding Gaussian noise:

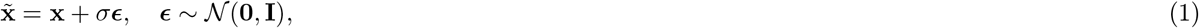

where the noise level *σ* ranges from near zero to large values, such that 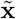 transitions from clean data to approximately pure noise. The reverse process is characterized by the score function 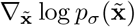, which guides denoising via a stochastic differential equation (SDE) or its deterministic probability flow ODE.

Following the EDM framework^57^, we parameterize the denoiser network with explicit preconditioning to ensure stable training dynamics. Specifically, the denoiser is defined as:

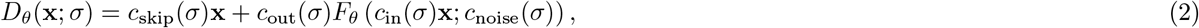

where *F*_*θ*_ denotes the neural network, and the preconditioning functions are designed to maintain unit variance for both network inputs and training targets:

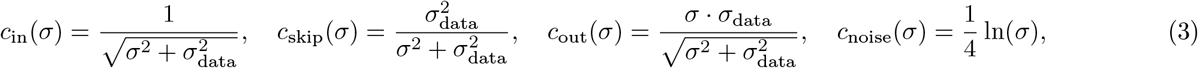

where *σ*_data_ is the standard deviation of the data distribution. *c*_skip_(*σ*) denotes the skip connection, while *c*_in_(*σ*) and *c*_out_(*σ*) scale the input and output magnitudes respectively.

The training objective of the EDM framework employs a weighted denoising loss:

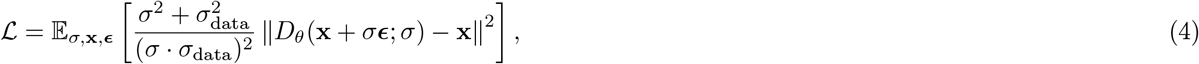

where the weight 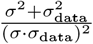 balances the effective loss across different noise levels. The noise level *σ* is sampled from a log-normal distribution ln 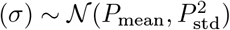, which concentrates training efforts on intermediate noise levels where the denoising task is both learnable and perceptually meaningful. We also add several auxiliary losses to ensure structural and distributional correctness of our generated trajectories. In particular, we propose a physically grounded dynamics loss that trains BioKinema to match the kinetic and thermodynamic behavior of the reference dynamics. The full details of our training objective are provided in the **Supplementary information**.

In our setting, each conformation **x**_*t*_ ∈ ℝ^*N* × 3^ represents a frame in a biomolecular trajectory, and diffusion models are used to denoise all frames in a trajectory.

### Model Architecture

Our model leverages the architectural foundation of AlphaFold 3 (AF3)^8^. The original AF3 framework is specially designed for static structure prediction. We extend this framework to capture the continuous temporal evolution required for molecular dynamics trajectory generation. As illustrated in **Figure 1**, the overall pipeline consists of three main components: an input embedder, a pairformer module, and a spatial-temporal diffusion module.

Given an input biomolecular system, the **Input Embedder** (3 blocks) first generates two types of representations: a single representation of dimension (*n, c*_*s*_) encoding per-token features, and a pair representation of dimension (*n, n, c*_*p*_) encoding pairwise relationships, where *n* is the number of tokens (polymer residues and ligand atoms). These representations are further processed by optional Template and MSA modules to incorporate structural and evolutionary information. The **Pairformer** module then serves as the dominant processing block that refines the pair and single representations through triangle updates and attention mechanisms. In our framework, the Input Embedder and Pairformer parameters are initialized from pre-trained Protenix^31^, the open-source reproduction of AlphaFold 3, and remain frozen during training. This design choice leverages the rich structural and evolutionary priors captured by AF3 and significantly reduces the training computational cost.

The refined pair and single representations, together with the input representations, are then passed to our **Spatial-Temporal Diffusion Module**, which introduces a physically motivated architecture to generate continuous all-atom MD trajectories. We describe the key designs in the following sections.

#### Spatial-Temporal Diffusion Module

The original AF3 diffusion module operates directly on raw atom coordinates; it is trained to receive noisy atomic coordinates and predict the true coordinates. The module consists of sequence-local attention blocks operating on atoms and global attention blocks operating on tokens. While this design is effective for single-structure prediction, it only models spatial relationships between atoms/tokens within a single frame and cannot capture temporal dependencies across trajectory frames.

To enable trajectory generation, we extend the attention mechanism to jointly model both spatial and temporal dimensions. Specifically, for each atom or token, we perform attention not only across different atoms/tokens within the same frame (spatial attention), but also across all frames for the same atom/token (temporal attention). This spatial-temporal joint modeling allows the network to learn how atomic positions evolve over time.

#### Physically Motivated Temporal Attention with Exponential Decay

A central question in designing temporal attention is how to encode the temporal relationships between frames. Standard positional encoding used in language models, such as sinusoidal embeddings^58^ or rotary position embeddings^59^, assumes discrete spaced positions and lacks a natural notion of physical time for molecular dynamics. We instead design our temporal attention bias based on the statistical properties of equilibrium MD trajectories.

#### Continuous-Time Attention Bias

We adopt an attention bias that is a function of the time difference between frames. Specifically, for frames *i* and *j* at times *t*_*i*_ and *t*_*j*_, we model the temporal self-attention matrix as:

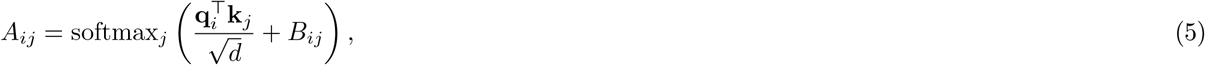

where the bias matrix *B* has entries:

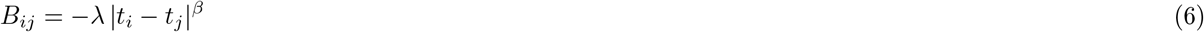

where *β* ≥ 0 is a stretching exponent, and *λ >* 0 is a learnable decay factor. Since the softmax function exponentiates its inputs, the bias translates to a multiplicative stretched-exponential factor in the attention weights:

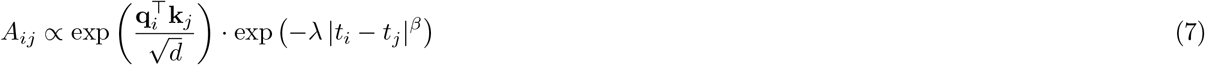

The attention weight between frames *i* and *j* thus combines content-based similarity (the 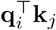 term) with a temporal proximity prior that decays monotonically with the time separation. The continuous form |*t*_*i*_ − *t*_*j*_ |^*β*^, in contrast to the integer-token distance used in standard ALiBi, lets the same model operate on trajectories with arbitrary, possibly non-uniform frame spacing.

#### Physical Motivation

The design of a temporal bias is motivated by the stochastic dynamics of molecular systems. In molecular dynamics simulations, the solvent environment acts as a thermal bath, introducing both systematic friction and random fluctuations that are naturally captured by Langevin dynamics^60^, which provides an effective description for the slow conformational coordinates. In aqueous solution, atoms lose their initial velocity due to solvent friction on a timescale of 10–100 femtoseconds^61^. This characteristic time, known as the momentum relaxation time, is negligible compared to the nanosecond-to-microsecond timescales of our training data. Consequently, inertial effects can be ignored, justifying the use of the overdamped (high-friction) limit^62, 63^. In this regime, the dynamics of a coordinate *x*(*t*) in a potential *V* (*x*) follows the stochastic differential equation:

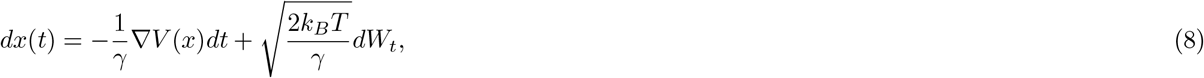

where *γ* is the friction coefficient, *k*_*B*_*T* is the thermal energy, and *W*_*t*_ denotes the standard Wiener process.

To analyze the local dynamics within a metastable state, we assume the potential energy surface can be approximated by a harmonic potential 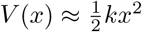 (via Taylor expansion around the equilibrium)^32^, where *k* is the force constant (stiffness) and *x* is the system coordinate. Substituting the harmonic force − Δ*V* (*x*) = − *kx* into Eq. 8 yields the Ornstein-Uhlenbeck (OU) process:

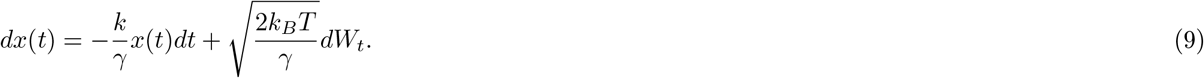

Defining the decay factor *λ* = *k*/*γ* and the noise amplitude 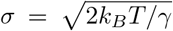, the explicit solution for the displacement is given by:

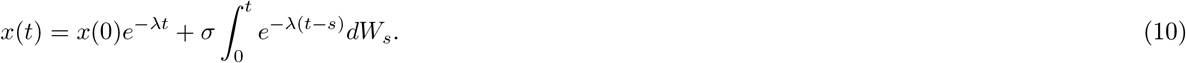

Note that in the integral term, *s* represents the integration time variable running from 0 to *t*.

A fundamental result for the stationary OU process is its auto-covariance function *K*(*τ*), which quantifies the correlation between the system’s coordinates at time *t* and *t* + *τ* ^60^:

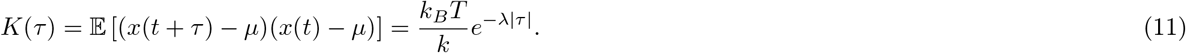

Here, *μ* = 0 is the equilibrium mean coordinate. This equation states that correlations decay exponentially with a characteristic timescale *τ*_*r*_ = 1/*λ*. Normalizing by the variance *K*(0) = *k*_*B*_*T*/*k* yields the autocorrelation function:

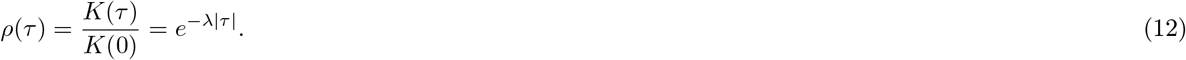

The OU analysis above yields a strictly linear bias, i.e. a pure exponential decay with *β* = 1. In practice, however, the physical time intervals spanned by our training data cover three to four orders of magnitude (from sub-nanosecond to microsecond scales). With a linear bias, a large time separation drives *B*_*ij*_ to a large negative value that, after the softmax, suppresses the attention weights of nearly all heads to zero; the model then loses access to distant frames, and the inter-frame correlations decay too rapidly to retain useful structural context. We therefore introduce the exponent *β* ∈ (0, 1] as an empirical correction to the OU form, replacing the exponential decay with a stretched-exponential decay. The sublinear growth of |*t*_*i*_ − *t*_*j*_|^*β*^ compresses this wide dynamic range, so that long-time biases remain informative rather than saturating.

#### Bidirectional Attention from Time-Reversal Symmetry

Equilibrium MD dynamics are characterized by time-reversal symmetry^64^. The joint distribution satisfies 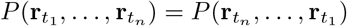, and correlations depend only on the time interval magnitude:

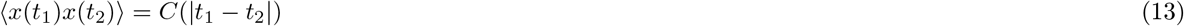

This implies that reversing the frame order produces a statistically indistinguishable trajectory, meaning information from future frames is equally valuable as information from past frames. A fundamental distinction arises when comparing this to text sequences: auto-regressive language models factorize the joint distribution as *P*(*w*_1_, …, *w*_*n*_) = Π_*i*_ *P*(*w*_*i*_|*w*_*<i*_), explicitly breaking time-reversal symmetry. Restricting to causal attention would therefore discard useful information without a physical justification.

Based on these considerations, we employ bidirectional attention with symmetric bias bias(*i, j*) = − *λ* |*t*_*i*_ − *t*_*j*_|^*β*^, respecting both the time-reversal symmetry and the stretched-exponential decay of temporal correlations in equilibrium dynamics.

#### Multi-Head Attention for Multiple Timescales

Biomolecular systems exhibit dynamics across a wide spectrum of timescales. For example, local fluctuations such as side chain rotamer transitions occur on the nanosecond scale^65^, whereas collective conformational changes, such as domain motions or local unfolding, occur over microsecond-to-millisecond timescales^65^. To capture this hierarchy, we allow each attention head *h* to learn a distinct decay factor *λ*_*h*_, and the final representation is a concatenation of features aggregated over different temporal horizons. Heads with large *λ*_*h*_ act as short-term memory filters focusing on fast, local dynamics, while heads with small *λ*_*h*_ serve as long-term memory filters capturing slow, global evolution. The decay factor *λ*_*h*_ for each attention head is a learnable parameter, and we initialize those decay factors following ALiBi^66^.

#### Continuous Time Support

Since the attention bias depends only on the time difference |*t*_*i*_− *t*_*j*_|^*β*^ rather than absolute timestamps, our model inherently supports arbitrary floating-point time intervals. During training, we randomly sample frame intervals ranging from 0.08 ns to 100 ns, enabling the model to learn dynamics across multiple timescales. At inference time, the time interval can be freely specified to generate trajectories at any desired temporal resolution.

#### Empirical Validation and Interpretation

To verify our assumption, we examined whether the temporal correlations in actual MD trajectories align with this exponential decay. As shown in **Supplementary Figure 15**, we first observe that the autocorrelation functions of C*α* coordinates exhibit a characteristic decay over time.

To quantitatively assess whether a multi-head attention mechanism with stretched-exponential decay biases can capture these correlation patterns, we fitted the empirical autocorrelation curves using a linear combination of stretched-exponential decay functions. We used four components, which is identical to the minimum number of temporal attention heads in our model. This exponential mixture fitting method achieves an average *R*^2^ of 0.83-0.88 for different *β* (**Supplementary Figure 15**), demonstrating high-fidelity fits for structural parameters. Collectively, this result confirms that the stretched-exponential decay form of our temporal attention bias is physically motivated and consistent with the statistical properties of MD trajectories.

### Noise-as-Masking Training Paradigm

Previous video generation^67^ and MD trajectory^27^ approaches mostly treat historical frames as conditioning input, first encoding them with a separate encoder and then injecting the information into the generative model via AdaLN layers. However, the conditioning encoder can compress historical information, potentially losing subtle but critical details necessary for accurate prediction. To overcome these limitations, we adopt a “noise-as-masking” strategy inspired by Diffusion Forcing^68, 69^, as illustrated in **Figure 1e**.

The core insight is to reframe the noise scale in diffusion training *σ* as a control over information visibility. A clean conditional frame at *σ* = 0 is fully visible (“unmasked”), while a fully noised frame at *σ* = *σ*_max_ is completely obscured (“masked”). During training, we leverage this by applying independent noise levels *σ*_*t*_ to each frame in a trajectory sequence: conditioning frames receive *σ*_*t*_ = 0 (clean), while target frames receive *σ*_*t*_ ∼ *p*(*σ*) sampled from the training noise distribution. This creates a single, unified input containing both clean conditioning frames and noisy target frames. Specifically, we randomly designate the first frame or the last frame as conditioning (unmasked), enabling the model to learn both temporal forecasting and interpolation capabilities simultaneously.

The entire sequence is then processed by the **Spatial-Temporal Diffusion Module**. Because the temporal transformer can attend to all frames simultaneously, it learns to use the clean “unmasked” frames as context to denoise and predict the “masked” frames, eliminating the need for separate encoders or injection layers. This unified framework makes our model highly versatile during inference: by simply controlling which frames are clean and which are noised, our model can seamlessly perform **forecasting** (conditioning on initial frames to predict future frames), **interpolation** (conditioning on boundary frames to generate intermediate frames), or any combination thereof—all within a single, unchanged architecture.

### Hierarchical Sampling Strategy

#### Challenges in Long Timescale Simulation

Generating long molecular dynamics trajectories presents a fundamental challenge: error accumulation in auto-regressive generation, which is necessitated by the prohibitive computational memory costs of generating long trajectories for complex biomolecular systems. When generating trajectories purely through sequential forecasting, small prediction errors at each step accumulate over time, leading to increasingly unrealistic conformations and eventual trajectory divergence. For example, generating a 1 *μ*s trajectory at 0.1 ns resolution requires 10,000 frames. If we generate in blocks of 50 frames (5 ns each), this demands 200 sequential forecasting steps, with errors accumulating at each iteration.

#### Forecasting + Interpolation Combined Strategy

To address this challenge, our hierarchical sampling strategy leverages both forecasting and interpolation capabilities enabled by the noise-as-masking paradigm. The key insight is that **forecasting determines the evolution of the global trajectory**, while **interpolation fills in local details**.

The two stages of our hierarchical framework are implemented simply by applying different masking and time schedules to our unified model during inference.

#### Coarse-grained Forecasting

The first stage generates a coarse-grained trajectory by sampling at large time intervals Δ*t*_coarse_, resulting in a sparse sequence **X**_*C*_ ={**x**_0_, **x**_*k*_, **x**_2*k*_, …}where *k* = Δ*t*_coarse_/Δ*t*_fine_. This stage establishes the overall trajectory evolution and captures slow, large-scale conformational changes.

This task is framed as a forecasting problem where, given the initial frame **x**_0_, the model generates all subsequent coarse-grained frames. During inference, this setup supports multiple generation strategies:

- **All-at-once:** All future frames {**x**_*k*_, **x**_2*k*_, …} are generated concurrently. The initial frame is provided as clean input, while all other frames are initialized from noise and denoised simultaneously using an SDE solver.
- **Auto-regressive (AR):** Frames are generated in sequential blocks of size *j*. To generate each block, the model conditions on previously generated frames (set as clean, “unmasked” inputs) and simultaneously denoises all frames within the current block. Once generated, this block is appended to the conditioning set, and the process repeats until the full coarse trajectory is complete.

By using large time intervals (e.g., Δ*t*_coarse_ = 5–20 ns), the coarse-grained stage requires far fewer forecasting steps. For the 1 *μ*s example above, using Δ*t*_coarse_ = 5 ns with 200 ns generation blocks requires only 5 forecasting iterations instead of 200. This reduction yields two key advantages: it significantly mitigates error accumulation by reducing sequential steps, and it expands the model’s temporal receptive field by generating longer trajectory segments, enabling better capture of global protein kinetics.

#### Fine-grained Interpolation

While the coarse-grained trajectory captures global dynamics, many applications require fine temporal resolution to observe detailed molecular events such as fast local fluctuations. Generating such fine-grained trajectories directly through forecasting would suffer from severe error accumulation.

Our framework addresses this through interpolation: after obtaining the coarse-grained trajectory {**x**_0_, **x**_*k*_, **x**_2*k*_, …}, the second stage fills in the intermediate frames. For each coarse interval, we generate the frames {**x**_*ik*+1_, …, **x**_(*i*+1)*k*−1_} conditioned on the two “anchor” frames **x**_*ik*_ and **x**_(*i*+1)*k*_.

During inference, interpolation is performed in an “all-at-once” manner. The anchor frames are provided as clean inputs (*σ* = 0), while all intermediate frames are initialized from noise (*σ* = *σ*_max_). The model then simultaneously generates all *k* − 1 intermediate frames through the standard denoising process. Crucially, because both endpoints are fixed, the interpolation task is geometrically constrained and does not suffer from unbounded error growth when compared to pure forecasting.

Collectively, this hierarchical approach of coarse forecasting followed by fine interpolation enables BioKinema to efficiently generate long, physically plausible trajectories at arbitrary temporal resolutions, combining the flexibility of forecasting with the stability of interpolation.

### Training Dataset

To let the model capture a diverse range of molecular dynamics across different timescales and system types, we curated a comprehensive training dataset comprising approximately 54,600 *μ*s (54.6 ms) of MD simulations (**Table 1**). The corpus combines three general-purpose MD repositories (ATLAS, MISATO, and MDposit) with three large-scale datasets from the BioEmu MD release^19^ that broaden conformational and sequence diversity:

**Table 1.** Summary of the molecular dynamics datasets used to train BioKinema. Counts reflect the systems and trajectories retained in the training indices after preprocessing. Force fields and temperatures for the three BioEmu-derived datasets (Octapeptides, CATH1, CATH2) are taken from the released per-system metadata; aggregate times are rounded.

| Dataset | Systems | Trajectories | Aggregate (μs) | Force field | Temp. (K) |
| --- | --- | --- | --- | --- | --- |
| ATLAS<sup>36</sup> | 1,389 | 4,167 | ∼417 | CHARMM36m | 300 |
| MISATO<sup>14</sup> | 14,628 | 14,628 | ∼117 | Amber | 300 |
| MDposit<sup>70</sup> | 3,271 | 3,271 | ∼1,470 | Various | 298–300 |
| Octapeptides<sup>19,63</sup> | 1,100 | 5,496 | ∼5,800 | ff99SB-ILDN | 300 |
| CATH1<sup>19,63</sup> | 50 | 1,248 | ∼5,400 | ff99SB-ILDN | 300 |
| CATH2<sup>19,72</sup> | 1,040 | 40,984 | ∼41,400 | ff99SB-ILDN | 300 |
| **Total** | **21,478** | **69,794** | **∼54,600** |  |  |

#### ATLAS Dataset

We utilize the ATLAS dataset^36^, which provides high-quality trajectories for protein monomers. The dataset contains 1,389 protein chains, curated based on the ECOD (Evolutionary Classification of Protein Domains) classification to remove structural redundancy. It covers 97 of the top 100 most common ECOD domains. Simulations were performed using GROMACS with the CHARMM36m force field and TIP3P water model in a physiological saline environment (150 mM NaCl). Each system includes 3 independent replicates of 100 ns each, totaling approximately 417 *μ*s.

#### MISATO Dataset

To capture protein-ligand interactions, we utilize the MISATO dataset^14^, whose structures originally come from PDBbind. It comprises approximately 16,000 protein-ligand complexes simulated using the Amber20 force field with explicit TIP3P solvent. Each complex was simulated for 10 ns. The first 2 ns were discarded as the equilibration phase, and the final 8 ns (100 frames) were retained for training. After chemical-validity filtering, 14,628 complexes are used, contributing approximately 117 *μ*s of data specifically focused on binding pocket dynamics and cryptic pocket opening.

#### MDposit Dataset

The largest portion of our training data comes from MDposit^70^, an open repository for molecular dynamics. We selected 3,271 systems encompassing a wide variety of biomolecular systems, including protein monomers and multimers, protein-nucleic acid complexes, and protein-ligand complexes. This dataset offers significant diversity in simulation duration, with trajectory lengths ranging from a minimum of 2.47 ns to a maximum of 5,350 ns, and an average length of 449.40 ns per trajectory, totaling 1,470 *μ*s.

#### Octapeptides Dataset

The octapeptides dataset consists of 1,100 unique eight-residue peptides^63^. Lewis et al.^19^ generated five 1 *μ*s trajectories per peptide (some extended to 2 *μ*s) using the Amber ff99SB-ILDN force field^71^ at 300 K. The processed set, saved at 10 ns per frame, comprises 5,496 trajectories and approximately 5,800 *μ*s of simulation.

#### CATH Dataset

We include two CATH-derived datasets from the BioEmu release, both simulated with the Amber ff99SB-ILDN force field^71^ at 300 K and saved at 10 ns per frame. **CATH1** consists of 50 CATH protein domains^63^; Lewis et al.^19^ extended the original simulations (four 0.5 *μ*s trajectories per domain) to a cumulative duration exceeding 100 *μ*s per domain; each domain has 23–25 trajectories (median 25) spanning 0.5–10 *μ*s (median 5 *μ*s), for a total of 1,248 trajectories and approximately 5,400 *μ*s. **CATH2** prioritizes sequence coverage: it includes 1,040 domains of 50–200 residues selected from the CATH database^72^, each sampled with roughly 40 trajectories of 1 *μ*s, giving 40,984 trajectories and approximately 41,400 *μ*s in total.

#### Data Preprocessing and Sampling

We applied a standardized preprocessing pipeline across all datasets. First, we adopt an implicit solvent representation following previous methods^8, 19, 27^ and removed all solvent molecules from the trajectories. Second, missing residues in the simulation topology were identified by aligning with the reference PDB structure; these residues were marked as “unresolved,” with their corresponding loss terms masked during training. Third, for systems containing small molecules, we performed valency checks and filtered out trajectories exhibiting chemical validity issues.

During training, we employed a dynamic sub-sampling strategy to construct training samples. For each trajectory, the inter-frame interval Δ*t* is drawn log-uniformly between the native frame spacing and up to roughly 25% of the trajectory length. The resulting effective intervals are approximately 1–25 ns for ATLAS, 0.08–2 ns for MISATO, and 0.1–100 ns for MDposit. Octapeptides, CATH1, and CATH2 are saved at 10 ns per frame, giving intervals from 10 ns up to 250 ns.

## Resource availability

All original code will be publicly available on GitHub https://github.com/IDEA-XL/BioKinema.

## Acknowledgments

We would like to express our gratitude to Dr. Han Ke and Dr. Zequn Liu from Peking University, and Dr. Yi Isaac Yang and Dr. Maodong Li from Shenzhen Bay Laboratory, for their valuable discussions and assistance with this work. We also thank Prof. Simon Olsson from Chalmers University of Technology for his insightful comments on the evaluation experiments.

This project was supported by Shenzhen Hetao Shenzhen-Hong Kong Science and Technology Innovation Cooperation Zone, under Grant No. HTHZQSWS-KCCYB-2023052.

## Supplementary information

### Training Objective

Our training objective supervises both the per-frame structural accuracy and the statistical and kinetic fidelity of the generated trajectories. It comprises four families of terms: (i) a per-frame structure reconstruction loss ℒ _struct_; (ii) ensemble flexibility losses ℒ _flex_ that reproduce the conformational fluctuations of the reference simulations; (iii) dynamics losses ℒ _dyn_ that enforce correct temporal correlations and state-to-state kinetics; and (iv), for ligand-unbinding fine-tuning, a ligand geometric center loss ℒ _center_. For standard MD-trajectory training, the total loss is

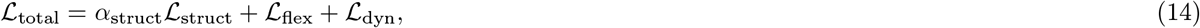

where ℒ _flex_ and ℒ _dyn_ are the weighted sums defined below. For unbinding trajectory prediction, the ensemble and dynamics terms are replaced by the ligand geometric center loss, ℒ _total_ = *α*_struct_ ℒ _struct_ + *α*_center_ ℒ _center_. All loss weights and the associated MSM/TICA hyperparameters are summarized in **Table 2**.

**Table 2.**
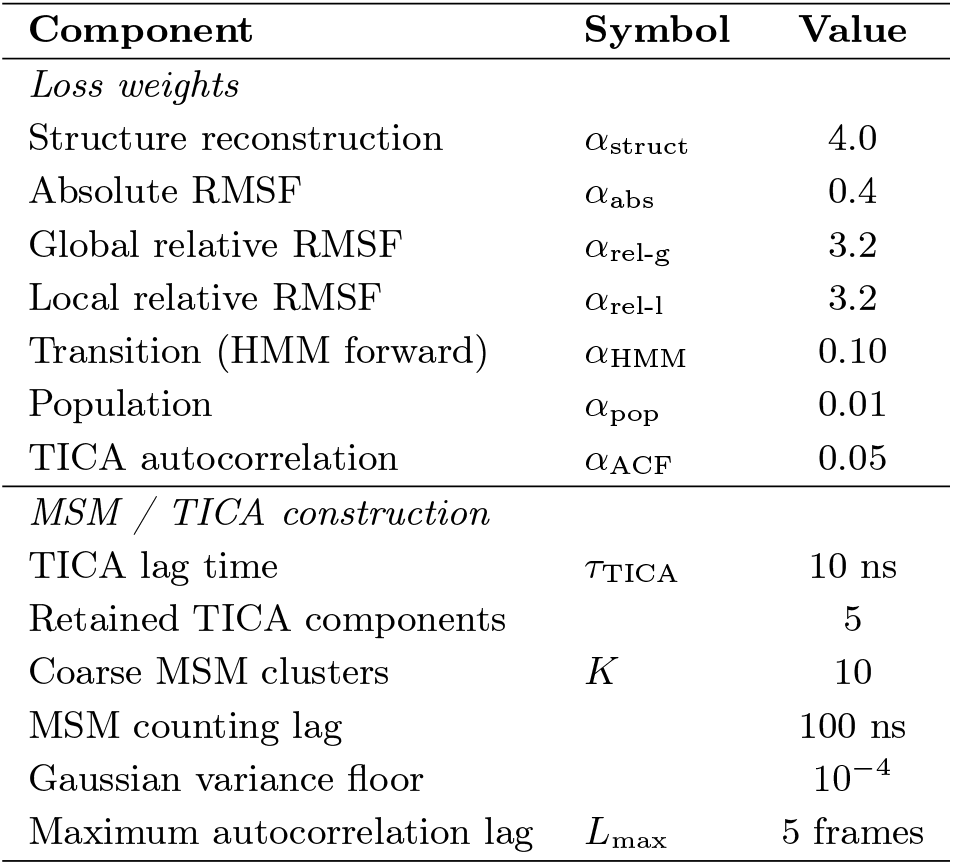
Loss weights and MSM/TICA settings for full-trajectory training. During unbinding fine-tuning, the ensemble and dynamics terms are replaced by the ligand geometric center loss.

| Component | Symbol | Value |
| --- | --- | --- |
| *Loss weights* |  |  |
| Structure reconstruction | $\alpha_{\text{struct}}$ | 4.0 |
| Absolute RMSF | $\alpha_{\text{abs}}$ | 0.4 |
| Global relative RMSF | $\alpha_{\text{rel-g}}$ | 3.2 |
| Local relative RMSF | $\alpha_{\text{rel-l}}$ | 3.2 |
| Transition (HMM forward) | $\alpha_{\text{HMM}}$ | 0.10 |
| Population | $\alpha_{\text{pop}}$ | 0.01 |
| TICA autocorrelation | $\alpha_{\text{ACF}}$ | 0.05 |
| *MSM / TICA construction* |  |  |
| TICA lag time | $\tau_{\text{TICA}}$ | 10 ns |
| Retained TICA components |  | 5 |
| Coarse MSM clusters | $K$ | 10 |
| MSM counting lag |  | 100 ns |
| Gaussian variance floor | | $10^{-4}$ |
| Maximum autocorrelation lag | $L_{\text{max}}$ | 5 frames |

**Table 3.**
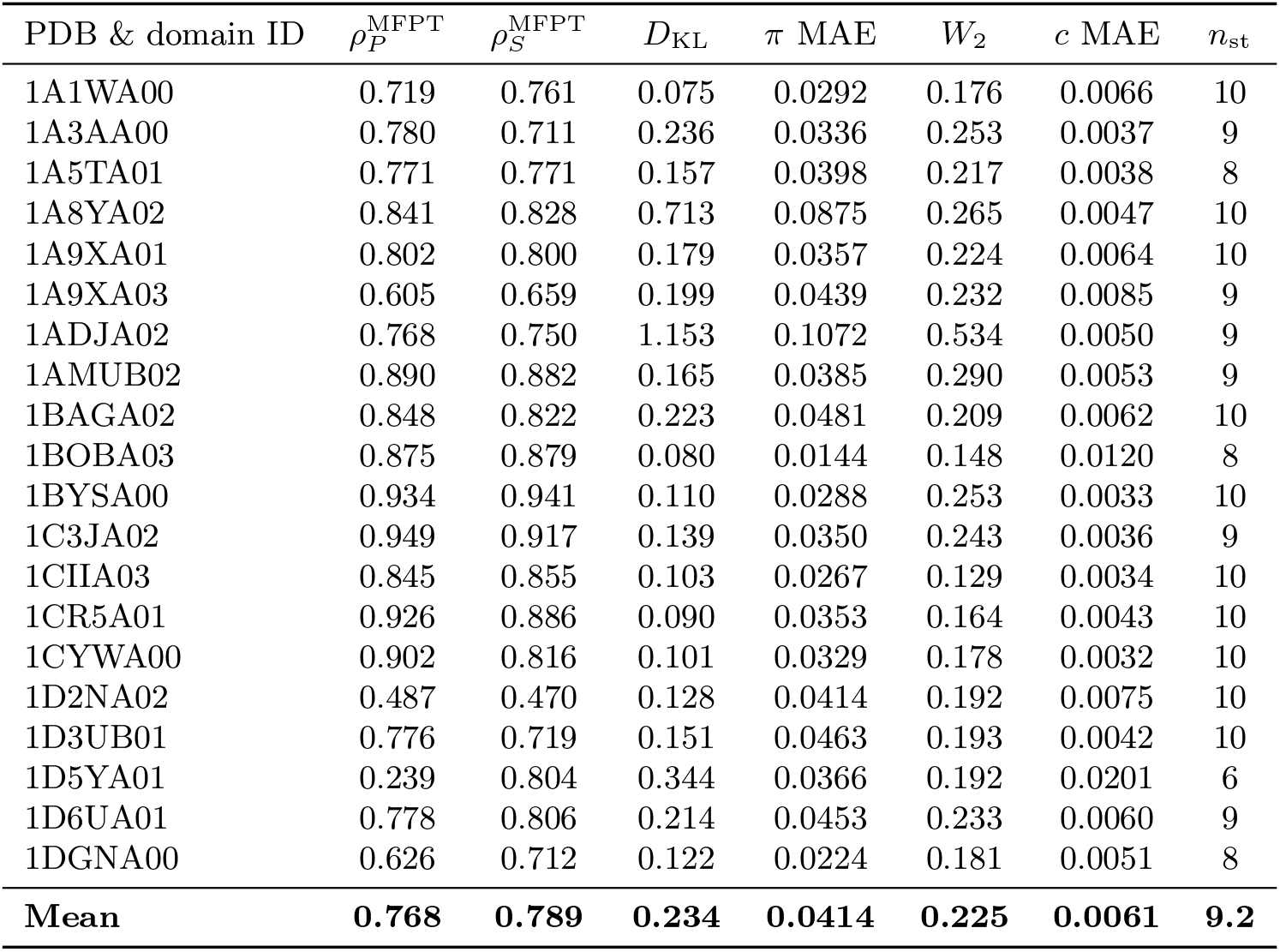
Per-system kinetics and stationary-distribution accuracy on the CATH2 test set. For each system: Pearson and Spearman correlation of log_10_(MFPT) between MD and BioKinema 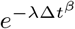, Kullback–Leibler divergence *D*_KL_(*π*_MD_ *π*_BK_) between the MD and BioKinema reversible-MLE MSM stationary distributions, the per-state mean absolute error (MAE) of *π*, the root-mean Wasserstein-2 (*W*_2_) distance across the five slowest TICA dimensions, the per-residue-pair C*α* contact-probability MAE between MD and BioKinema (*c* MAE; 8 Å contact cutoff), and the number of MSM states populated by BioKinema (*n*_st_, out of *K*=10; a state is counted as populated if *π*_BK_ *>* 10^−4^). The bottom row gives cross-system means.

#### Structure Reconstruction Loss

The structure reconstruction loss supervises the accuracy of individual trajectory frames produced by the diffusion module. Following AlphaFold 3^8^, we compute the loss on the fully denoised atomic coordinates output by the diffusion module.

##### Coordinate Supervision

Since the trajectory is treated as a unified entity, we aligned the training MD trajectories to the first frame to remove internal rigid-body motions and subsequently augmented them via global random rotations and translations. Consequently, the model is trained to predict coordinates in this global frame directly, and the loss is computed without further alignment:

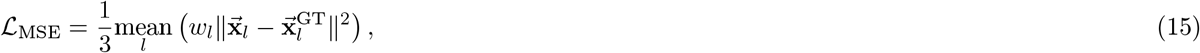

where the factor 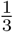 normalizes over spatial dimensions. The per-atom weights *w*_*l*_ upweight nucleotide and ligand atoms:

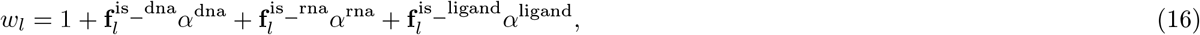

with binary indicators 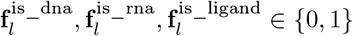 and hyperparameters *α*^dna^ = *α*^rna^ = 5, *α*^ligand^ = 10 following AlphaFold 3^8^.

##### Auxiliary Loss Components

Following AlphaFold 3^8^, we incorporate two auxiliary loss terms to enhance structural validity. First, a bond length loss ℒ _bond_ is applied to covalent bonds connecting ligands to protein chains, ensuring that these attachments maintain physically plausible geometries. Second, we utilize a smooth lDDT loss ℒ _smooth_lddt_, which serves as a differentiable approximation of the Local Distance Difference Test metric to supervise the accuracy of local atomic neighborhoods.

##### Combined Loss

The complete structure reconstruction loss is modulated by a noise-level-dependent weighting following the EDM framework^57^. This weighting reduces the loss contribution at high noise levels where accurate denoising is inherently difficult:

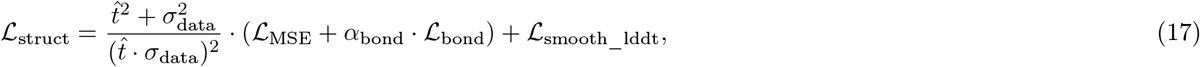

where 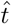 denotes the sampled noise level in the diffusion process, *σ*_data_ = 16 is a data-dependent constant, and *α*_bond_ = 1, all following AlphaFold 3^8^.

#### Flexibility Loss

We introduce flexibility losses that supervise the distribution of structural fluctuations rather than individual conformations. These losses operate on ensembles of predicted conformations 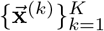 sampled from the model and corresponding reference MD trajectory frames 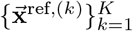, where the comparison is distributional rather than based on one-to-one frame correspondence. Consistent with the structure reconstruction objective, all reference MD trajectory frames are pre-aligned to the first frame to remove rigid-body motion before computing flexibility metrics.

##### Absolute Flexibility Loss

This loss encourages the model to reproduce per-atom positional fluctuations. For each atom *l*, we compute the root-mean-square fluctuation (RMSF) around the ensemble mean:

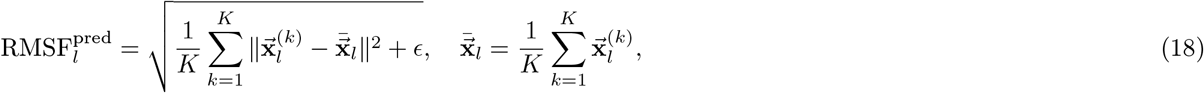

where *ϵ* is a small constant for numerical stability. The reference MD RMSF is computed analogously. The absolute flexibility loss is computed as follows:

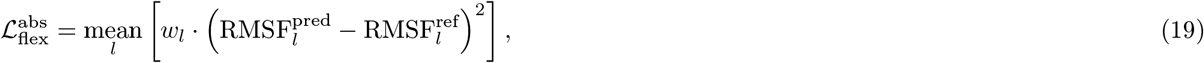

where *w*_*l*_ follows the same weighting scheme as in the structure reconstruction loss.

##### Relative Flexibility Loss

While absolute flexibility captures global positional variance, relative flexibility characterizes distance fluctuations between atom pairs, which are invariant to rigid-body motions. We define two variants based on the type of atom pairs considered.

Global relative flexibility considers pairwise distances among functionally important atoms: C*α* for protein residues and all heavy atoms for ligands. For each qualifying atom pair (*l, m*), we compute the standard deviation of their pairwise distances across the ensemble:

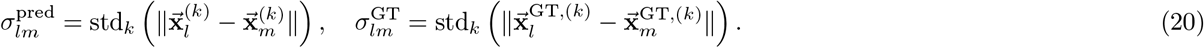

The global relative flexibility loss applies interaction-type-dependent weights *ω*_*lm*_ and distance-dependent weights *γ*_*lm*_.

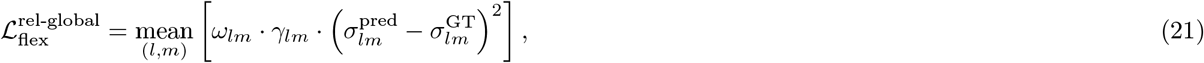

where *ω*_*lm*_ assigns higher weights to ligand-ligand pairs (*ω* = 10) and protein-ligand pairs (*ω* = 5) compared to protein-protein pairs (*ω* = 1), reflecting the importance of accurate ligand flexibility modeling. The distance weight *γ*_*lm*_ emphasizes proximal contacts: pairs with mean distance ≤ 5 Å receive weight 4, those within (5, 10] Å receive weight 2, and those *>* 10 Å receive weight 1.

Local relative flexibility focuses on intra-residue atom pairs, capturing side chain and local backbone fluctuations.

For atom pairs (*l, m*) belonging to the same residue (identified by shared token indices), we compute:

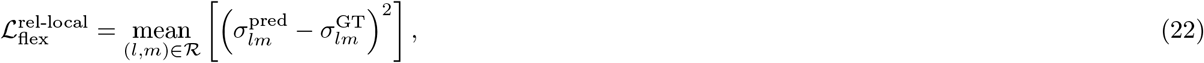

where ℛ denotes the set of intra-residue atom pairs.

##### Combined Flexibility Loss

The complete flexibility loss combines the three components,

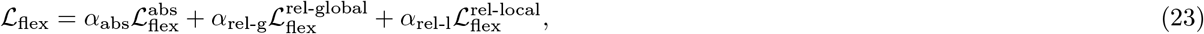

whose weights (Table 2) emphasize the relative, rigid-body-invariant pairwise-distance terms over the absolute RMSF term.

#### Dynamics Loss

We propose a physically grounded dynamics loss that uses a per-system Markov State Model (MSM) to supervise two physical properties of the generated trajectories: the kinetics, which correspond to the transition rates and relaxation timescales between metastable states, and the thermodynamics, which corresponds to the equilibrium populations of these states. The loss comprises three terms that share a common differentiable pipeline:

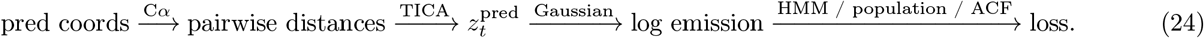

Here each frame *t* is projected into the five-dimensional TICA space, yielding the coordinate *z*_*t*_ ∈ *ℝ*^5^ (written 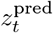 when distinguishing a model-generated frame from the ground truth); the three losses are defined below.

##### MSM Construction

For each system, we extract C*α* coordinates from all frames and use the upper-triangular C*α*–C*α* pairwise distances as structural features, which are invariant to global rotation and translation. We apply Time-lagged Independent Component Analysis (TICA)^40, 41^ to these features, fitting the TICA model on all available trajectories of the system, with a TICA lag time of 10 ns, and retain the five slowest independent components. The resulting 5-dimensional coordinates are clustered with K-means into *K* = 10 coarse metastable states, which define the MSM used by the dynamics losses. Because a reliable TICA/MSM model requires broad sampling of the conformational space, this dynamics-loss construction is applied only to the octapeptides and CATH datasets which can provide many replicas per system; systems with too few trajectories are therefore excluded from the dynamics losses.

We then estimate a reversible MSM. To capture slow dynamics reliably, the transition count matrix is estimated at a counting lag of 100 ns (10 frames). For compatibility with the per-frame (10 ns) time resolution used during training, we store the transition matrix at the per-frame lag, 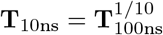, computed via eigendecomposition; this captures slow modes far more accurately than estimating directly at a 10 ns lag. The stationary distribution ***π*** is obtained by power iteration and is preserved under the matrix root.

For each coarse state *i*, we fit a diagonal Gaussian 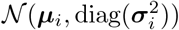 to the TICA coordinates of its frames, using the empirical mean and variance with a floor of 10^−4^. We use diagonal rather than full covariance so that out-of-distribution conformations visited during training still receive a non-negligible emission probability, preventing the loss from saturating and the gradients from vanishing.

##### Transition Loss (HMM Forward)

The transition loss evaluates how likely the generated trajectory is under the MSM, treating the trajectory as an observed sequence in a Hidden Markov Model with:

- Hidden states *s*_*t*_ ∈ {0, …, *K* − 1} — the MSM coarse states (*K* = 10).
- Initial distribution ***π*** — the MSM stationary distribution.
- Transition probabilities **T**^*k*^ — the stored 10 ns transition matrix raised to the power *k*, where *k* is the ratio of the actual frame spacing to the stored 10 ns lag, so that a frame pair separated by a physical interval of *k* · 10 ns is matched to **T**^*k*^. During training, *k* is sampled log-uniformly within [1, 25].
- Emission 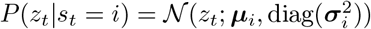— the per-cluster diagonal Gaussian from MSM construction.

The loss is the negative log-likelihood of the generated trajectory under this HMM, computed exactly via the forward algorithm (with *N*_frame_ the number of frames per trajectory) in *O*(*N*_frame_ · *K*^2^). The forward variable *α*_*t*_[*j*] = *P*(*z*_0_, …, *z*_*t*_, *s*_*t*_ = *j*) is the joint probability of the observed TICA coordinates of the first *t*+1 frames and the trajectory occupying state *j* at frame *t*; we evaluate it in log-space for numerical stability. The three lines below give, in order, (i) its *initialization* at the first frame as the stationary prior times the emission likelihood, (ii) its *recursion*, in which *α*_*t*+1_[*j*] accumulates the emission at frame *t*+1 and marginalizes over the previous state *i* weighted by the transition probability 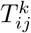, and (iii) the final *loss*, the length-normalized negative log-likelihood obtained by summing *α* over the last frame’s state:

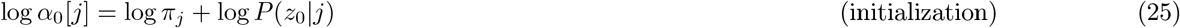

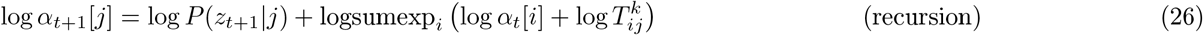

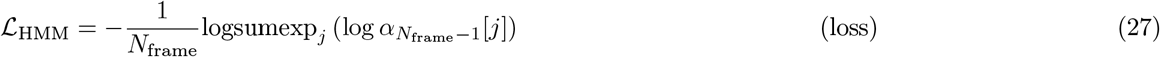

This loss captures both the thermodynamic correctness (via emission match to metastable states) and the kinetic correctness (via transition probabilities between states) in a single unified objective. The forward algorithm marginalizes over all 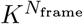 possible state paths exactly, avoiding the independence assumption of simpler state-assignment approaches.

##### Population Loss

The population loss ensures that the time-averaged state occupancy of the generated trajectory matches the trajectory’s empirical state distribution. Given the log emission probabilities, we compute a Bayesian soft assignment using the MSM stationary distribution ***π*** as a prior:

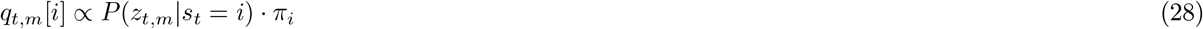

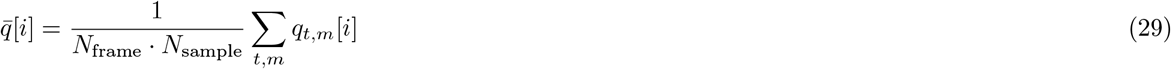

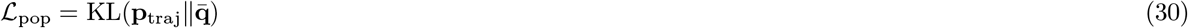

where *q*_*t,m*_[*i*] is the posterior probability that frame *t* of generated sample *m* occupies state *i, N*_sample_ is the number of generated samples, and **p**_traj_ is the empirical state distribution of the ground-truth trajectory; the MSM stationary distribution ***π*** serves as a fixed prior for the soft assignment.

##### TICA Autocorrelation Loss

The TICA autocorrelation loss enforces that the generated trajectory reproduces the correct relaxation timescales of the protein’s slowest motions. In the TICA basis, each component *d* decorrelates exponentially with a characteristic rate given by the TICA eigenvalue *λ*_*d*_:

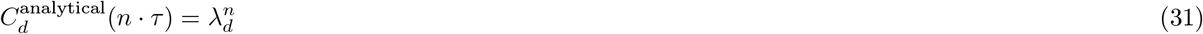

where *τ* is the TICA lag time and *n* is the number of lag steps. We compare this analytical prediction against the empirical autocorrelation computed from the generated trajectory:

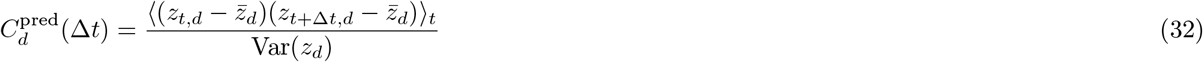

The loss is the MSE between empirical and analytical autocorrelation across all TICA dimensions and lag steps up to a maximum lag *L*_max_:

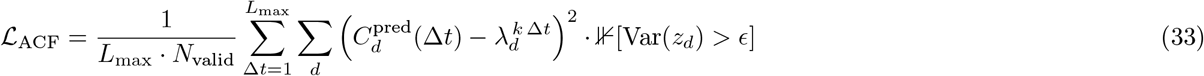

where *k* is the same frame-to-lag ratio defined above, so that the frame lag Δ*t* corresponds to *k* Δ*t* TICA lag steps. TICA dimensions with near-zero variance in the predicted trajectory are masked out by the indicator ⊮[] (threshold *ϵ* = 10^−3^) to prevent numerical blow-up from collapsed modes, and *N*_valid_ counts the surviving (dimension, lag) pairs over which the mean is taken.

##### Combined Dynamics Loss

Collecting the coordinate-space and TICA-space terms, the total dynamics loss is

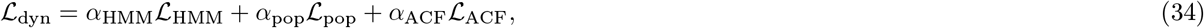

with weights given in Table 2. The whole pipeline is differentiable from C*α* coordinates through pairwise distances, the TICA projection, and the Gaussian emission model, enabling end-to-end gradient flow from the dynamics objectives back to the diffusion-model parameters.

#### Ligand Geometric Center Loss

For unbinding trajectory prediction, we replace the ensemble flexibility and dynamics losses with a ligand geometric center loss. Let ℐ_ligand_ denote the set of indices corresponding to ligand atoms, and *N*_*ℓ*_ =|ℐ_ligand_|be the number of ligand atoms. The geometric center at frame *t* is defined as:

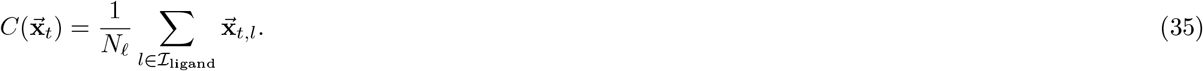

The loss penalizes discrepancies between the ligand centers of the predicted coordinates 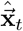 and the corresponding reference metadynamics coordinates 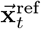, averaged over all trajectory frames:

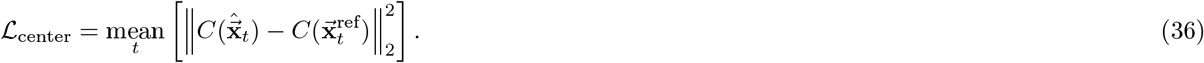

This term complements the structure reconstruction loss by softly anchoring the ligand’s global position during unbinding simulations, preventing spurious rigid translations while allowing the model to learn the dissociation pathway.

## Training and Inference Details

### Initialization and Architecture

To leverage the structural understanding of state-of-the-art static structure predictors, we initialize the Input Embedder and Pairformer modules with pre-trained weights from Protenix^31^, the open-source reproduction of AlphaFold 3. These modules remain frozen during training. To accelerate training throughput, we precompute the embeddings (single representation, pair representation, and single input) for all training data.

In the Spatial-Temporal Diffusion Module, the spatial attention and diffusion conditioning blocks are initialized from Protenix. The temporal attention layers are randomly initialized, with the exception of the output linear projection layers, which are initialized to zero. This “zero-initialization” strategy ensures that at the start of training, the model behaves identically to the single-frame AF3 predictor, allowing it to gradually learn temporal dependencies without disrupting the pre-trained spatial features.

### Temporal Attention Bias

We implement the temporal attention mechanism with multi-head attention. We set the number of attention heads to 4 for atom-wise temporal attention and 16 for token-wise temporal attention. We introduce a learnable bias decay factor *λ* in the continuous-time bias *B*_*ij*_ =− *λ* | *t*_*i*_ − *t*_*j*_|^*β*^, which differs across heads. These decay factors are initialized according to a geometric sequence ranging from 0.004 to 0.7 following ALiBi^66^, ensuring a broad coverage of temporal receptive fields. Specifically, larger *λ* values correspond to heads that focus on local, high-frequency fluctuations (e.g., side chain rotations), while smaller *λ* values allow heads to capture long-range, slow conformational changes (e.g., domain motions).

### Model Variants and the Choice of Stretching Exponent

Because the accessible timescales differ markedly across our data sources, we train two main variants of BioKinema. A *complex* variant, targeting protein–ligand complexes and short-timescale motions, is trained on ATLAS, MISATO, and MDposit, for which 97.4% of the trajectories span 8–100 ns; a *single-chain* variant, targeting long-timescale single-chain protein dynamics, is trained on Octapeptides and CATH, covering trajectory lengths of 0.5 to 10 *μ*s. For each variant we adopt the stretching exponent *β* that performs best on held-out data: *β* = 0.5 for the complex variant (selected on the ATLAS and MISATO tests; **Supplementary Figure 3**) and *β* = 0.25 for the single-chain variant (selected on the thermodynamic and kinetic tests; **Supplementary Figure 12**). Starting from BioKinema, we additionally fine-tune an unbinding variant on the DD-13M metadynamics data, for which *β* = 1 is used. We attribute this preference to the dominant timescale of each dataset: the single-chain variant spans the longest intervals (0.5–10 *μ*s), the complex variant intermediate ones (8–100 ns), and the unbinding variant the shortest (mean trajectory length of only 48 ps). A larger *β* drives the bias of widely separated frame pairs to large negative values and suppresses most attention heads, whereas a smaller *β* leaves frames at different separations poorly distinguished; the optimal *β* therefore decreases as the characteristic timescale of the data grows.

Unless otherwise noted, the main-text experiments use these variants as follows: the complex variant is used for the physical-stability and flexibility experiments (**Figure 2**) and the protein–ligand interaction experiments (**Figure 4**); the single-chain variant is used for the kinetics and thermodynamics experiments (**Figure 3**); and the unbinding variant is used for the ligand-unbinding experiments (**Figure 5)**.

### Training Hyperparameters

The model is trained using the Adam optimizer with a fixed learning rate of 10^−4^ and a linear warmup of 200 steps. We use a batch size of 32 distributed across 8 NVIDIA RTX A6000 GPUs. To balance the contribution of different data sources, we apply resampling within each epoch. For the complex variant, we resample the ATLAS, MISATO, and MDposit datasets to an approximate 1:1:1 ratio. For the single-chain variant, the CATH2, CATH1, and octapeptides datasets are sampled with relative weights of 0.444, 0.056, and 0.087, respectively. Following AF3, we apply random rotation and translation augmentation to the entire trajectory during training.

### Inference Configuration

We employ the EDM sampling framework for trajectory generation. Since the generation is conditioned on a known initial structure (Frame 0), the reverse diffusion process starts with strong structural guidance. Consequently, we find that a short rollout is sufficient for high-quality generation. We use a standard sampler with *N* = 20 steps, a noise scale parameter *σ*_noise_ = 1.75, and a step scale parameter *η* = 1.5.

### Evaluation metrics

#### Root Mean Square Deviation (RMSD)

RMSD measures the structural deviation between two conformations after optimal rigid-body alignment and is widely used to evaluate the overall stability of molecular trajectories. Given two structures containing (N) atoms, the RMSD is defined as

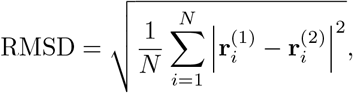

where 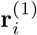 and 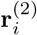 denote the Cartesian coordinates of atom (i) in the two aligned conformations. For physical stability evaluation, RMSD was computed between generated conformations before and after energy minimization using the Amber99SB+GAFF2 force field. Lower RMSD values indicate that only minor structural corrections are required during relaxation and therefore suggest higher physical plausibility.

#### Root Mean Square Fluctuation (RMSF)

RMSF quantifies the positional flexibility of individual residues throughout a trajectory and is commonly used to characterize equilibrium dynamics. For residue *i*, RMSF is calculated as

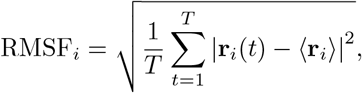

where **r**_*i*_(*t*) is the position of the residue (represented by its *C*_*α*_ atom) at time *t*, and ⟨**r**_*i*_⟩ denotes its time-averaged position over the trajectory. To evaluate the agreement between generated and reference trajectories, we computed the Pearson correlation coefficient between the per-residue RMSF profiles. High correlation indicates accurate reproduction of residue-level flexibility patterns.

#### Interaction Map Similarity

To assess whether BioKinema accurately reproduces protein–ligand interactions, we compared protein–ligand contact probability maps derived from generated trajectories and reference MD simulations. For each protein residue *i* and ligand heavy atom *j*, the contact probability is defined as

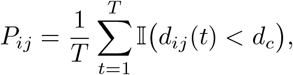

where *d*_*ij*_(*t*) is the distance between residue *i* and ligand atom *j* at frame *t, d*_*c*_ is the contact cutoff distance, and *I*(·) is the indicator function. We report interaction maps at two cutoffs: *d*_*c*_ = 5.0 Å, which captures all protein–ligand contacts, and *d*_*c*_ = 3.5 Å, which isolates strong, close-range interactions.

The interaction map similarity between generated and reference trajectories is then computed using the cosine similarity between the flattened contact probability matrices. Values closer to 1 indicate stronger agreement in the interaction landscape.

#### Mean First-Passage Time (MFPT)

MFPT is a fundamental kinetic observable that characterizes the expected time required for a system to reach a target state for the first time. After projecting trajectories into the time-lagged independent component (TICA) space and discretizing conformations into the *K*=10 Markov state model (MSM) states, MFPTs were computed from the MSM transition matrix.

For a transition from state *i* to state *j*, the MFPT *m*_*ij*_ satisfies

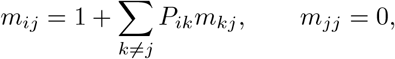

where *P*_*ik*_ denotes the row-stochastic transition probability from state *i* to state *k*. Because the MFPTs are computed from the stored per-frame transition matrix **T**_10ns_, solving this linear system yields *m*_*ij*_ in units of the 10 ns per-frame lag, which we multiply by 10 ns to obtain MFPTs in physical time. Note that this per-frame matrix is the 1/10 power of the count matrix estimated at the 100 ns counting lag (see MSM Construction above), so the 10 ns factor refers to the per-frame representation rather than the estimation lag.

To evaluate kinetic fidelity, pairwise MFPTs were computed for BioKinema and MD from their respective transition matrices on the shared set of *K*=10 MSM states, and compared over the state pairs reachable in both ensembles. Because MFPTs span several orders of magnitude, agreement was quantified by the Pearson correlation of log_10_(MFPT) together with the rank-based Spearman correlation. Accurate reproduction of MFPTs indicates that the generated trajectories preserve the relative ordering and timescales of conformational transitions.

### Experiment on Ligand Unbinding

Standard MD datasets predominantly capture local equilibrium fluctuations near a stable ground state. To evaluate our model’s capability to model non-equilibrium kinetics, we conducted specific experiments on ligand unbinding, which requires the model to predict the continuous trajectory of a ligand escaping the binding pocket. We fine-tuned BioKinema on the DD-13M dataset^17^, a large-scale metadynamics database specifically designed to capture ligand dissociation kinetics. DD-13M contains 26,612 dissociation trajectories across 565 complexes, recording the complete pathway from the binding state (L-P) to the unbinding state (L + P), each with an average of 48 ps length. In these metadynamics simulations, unbinding is determined by whether the ligand’s center of mass crosses the protein’s solvent-accessible surface; once the ligand’s center reaches this boundary, the simulation is terminated.

The metadynamics simulations in DD-13M use the ligand’s Center of Mass (COM) coordinates **R**_com_ = (*x, y, z*) as collective variables. A history-dependent bias potential *V* (*s, t*) is applied to these variables to discourage revisiting explored states and to force the ligand out of the binding pocket. Because the potential energy surface in metadynamics depends on the accumulated history of the simulation path, the underlying physics is inherently causal. Considering this, we apply a **causal mask** to the temporal attention mechanism during training or fine-tuning on the DD-13M dataset. This ensures that the prediction of a frame at time *t* depends only on the trajectory history *t*^*′*^ ≤ *t*, mirroring the accumulation of the bias potential. Moreover, because DD-13M trajectories are accelerated by metadynamics, their nominal time does not correspond to physical (unbiased) MD time. To bring the per-frame interval into the range the model has learned from equilibrium data, we scale the inter-frame time interval by a factor of 1000 when fine-tuning on DD-13M.

For evaluation, we define a challenging test set comprising 31 protein-ligand systems from DD-13M that share a maximum sequence identity of ≤ 40% with the training set. For evaluation, we generate 20 independent unbinding trajectories (60 ps) for each test system to observe the ligand’s escape from the binding pocket.

We evaluated the correctness of the generated trajectories and their ability to recover the dissociation pathways discovered by metadynamics. Since a given protein-ligand complex may have multiple valid exit routes, we computed precision and recall metrics by measuring whether the endpoints of generated trajectories correspond to any of the metadynamics-discovered exit points. For Metadynamics, the endpoint is originally defined as the point where the ligand centroid crosses the protein surface^17^; for our generated trajectories, it is defined as the point in the trajectory where the ligand is closest to the protein surface. Let ℰ = {**e**_1_, **e**_2_, …, **e**_*M*_}denote the set of *M* exit points observed in the metadynamics reference trajectories for a given complex, where each **e**_*j*_ represents the ligand centroid position in the final frame of a reference unbinding trajectory. Similarly, let *G* = {**g**_1_, **g**_2_, …, **g**_*N*_} denote the set of *N* generated trajectory endpoints for the same complex. Given a distance threshold *τ*, we define:

- **Precision**. This metric measures what fraction of generated trajectories terminate near a known exit point. A generated endpoint **g**_*i*_ is considered a match if it falls within distance *τ* of any reference exit point:

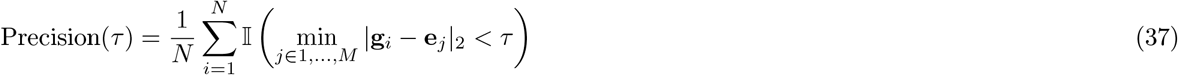

where *I*() is the indicator function. Higher precision indicates that generated trajectories reliably find valid exit routes.
- **Recall**. This metric measures what fraction of the known exit pathways are recovered by the generated trajectories. A reference exit point **e**_*j*_ is considered covered if at least one generated endpoint falls within distance *τ* :

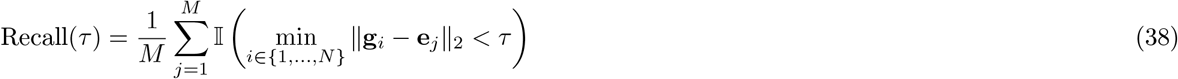

Higher recall indicates broader coverage of the diverse dissociation routes discovered by metadynamics.

#### Algorithm 1 Hierarchical Trajectory Generation (Main Pipeline)

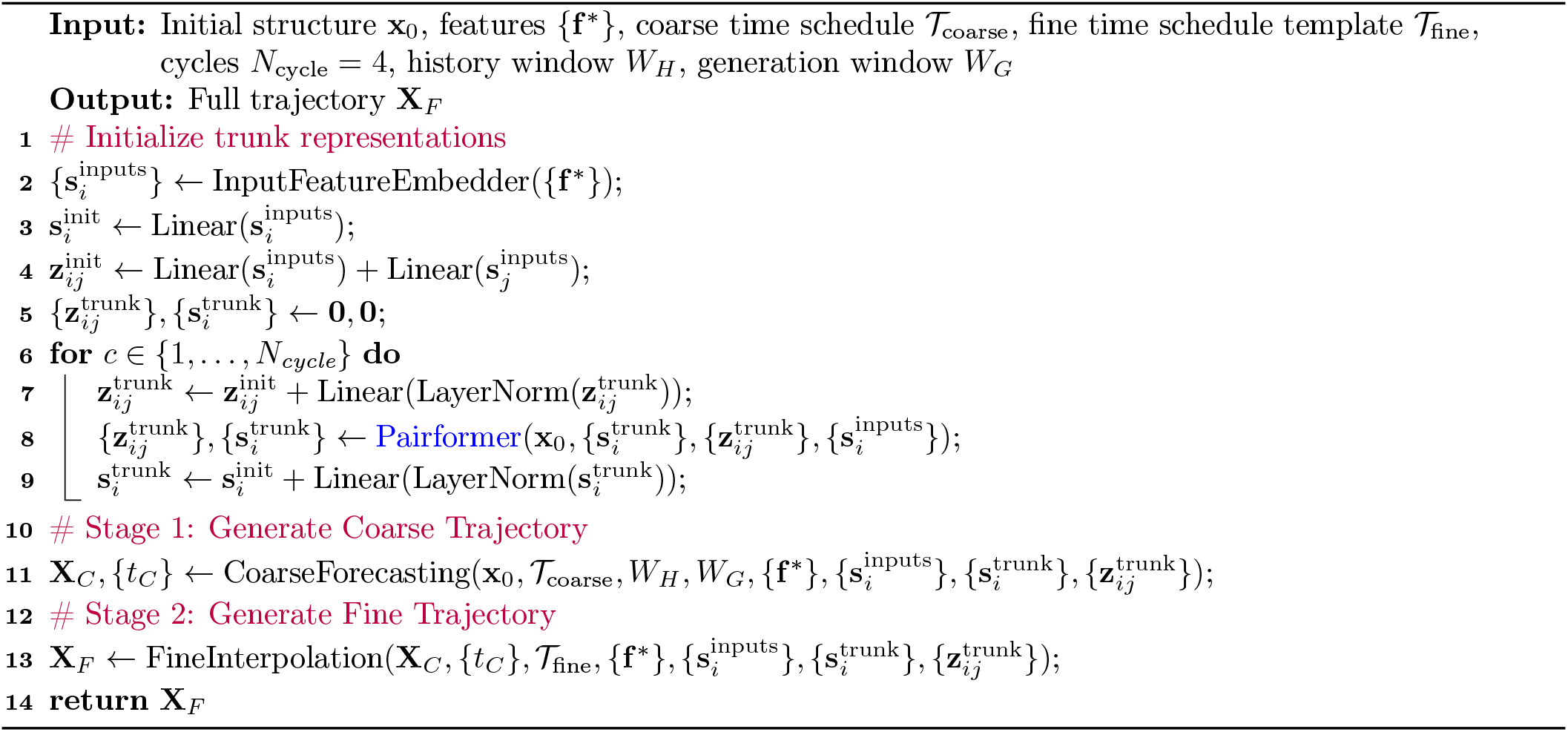

#### Algorithm 2 Stage 1: Coarse-grained Forecasting

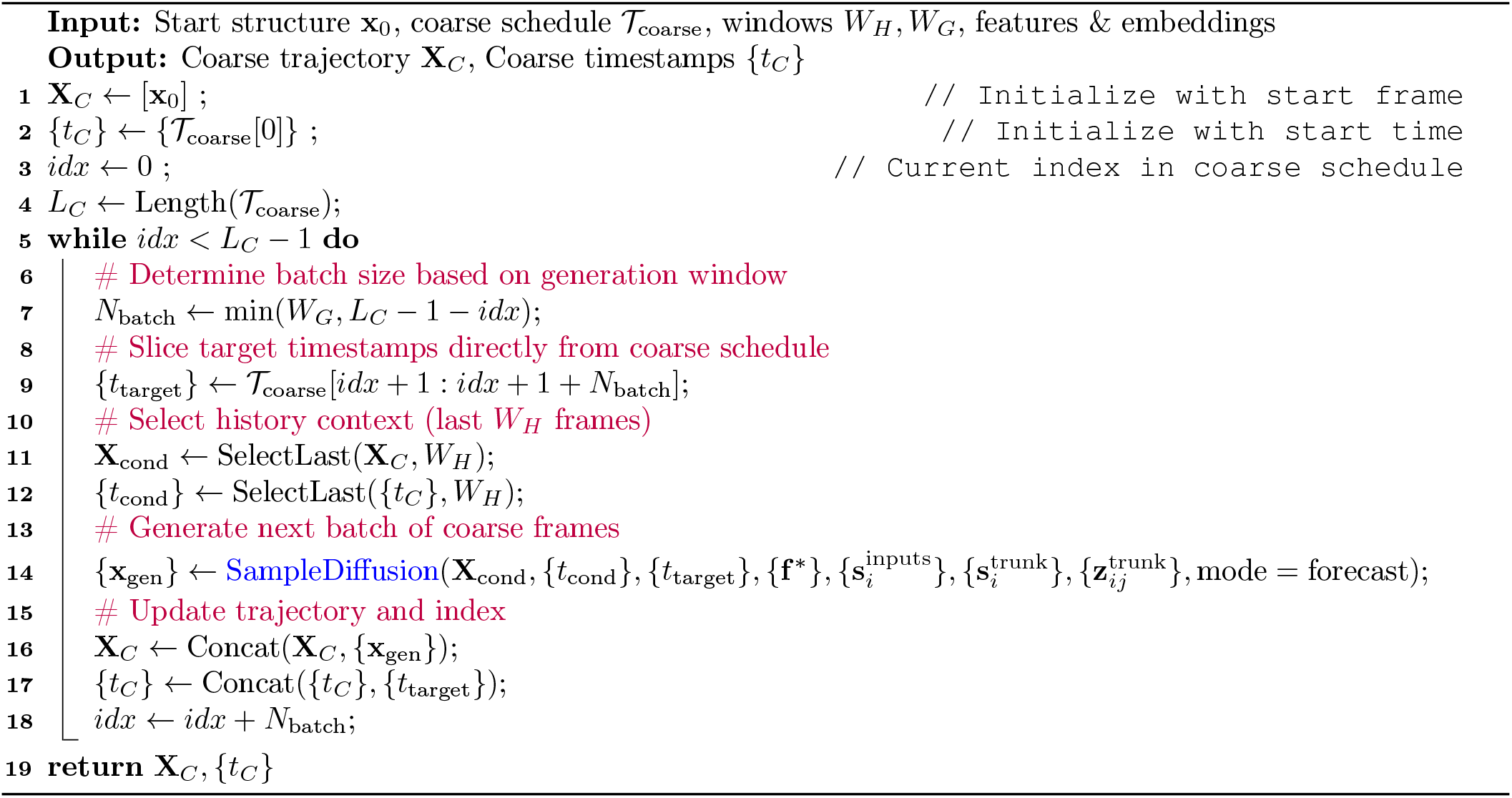

#### Algorithm 3 Stage 2: Fine-grained Interpolation

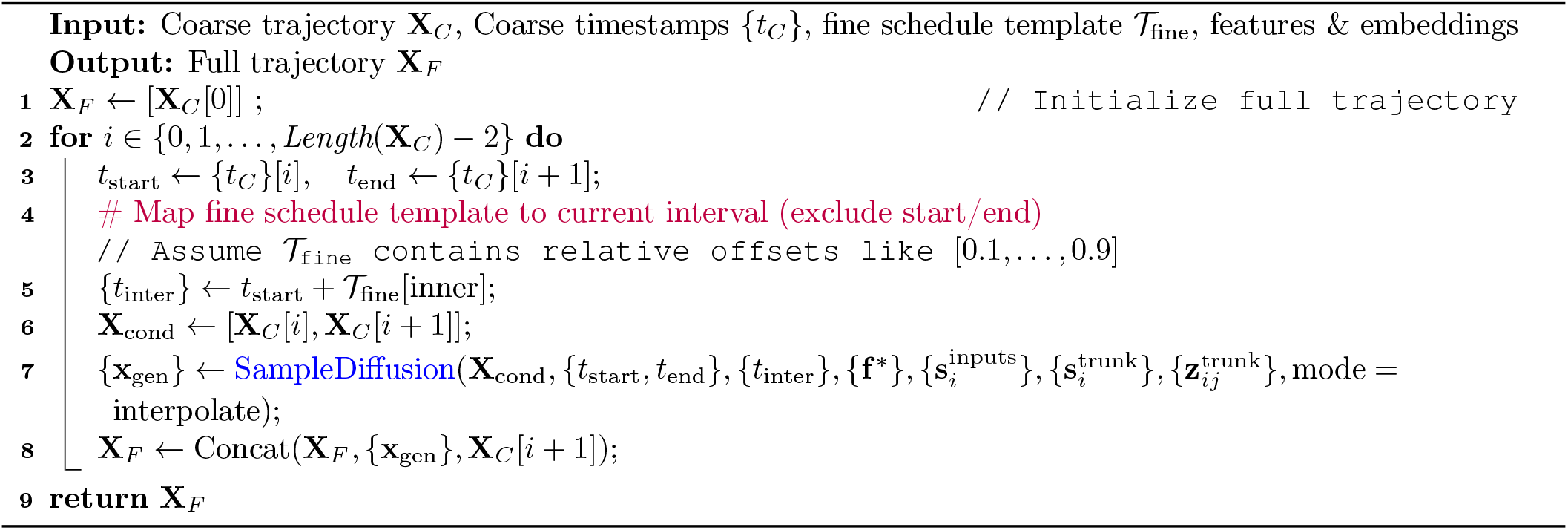

#### Algorithm 4 Training with Noise-as-Masking

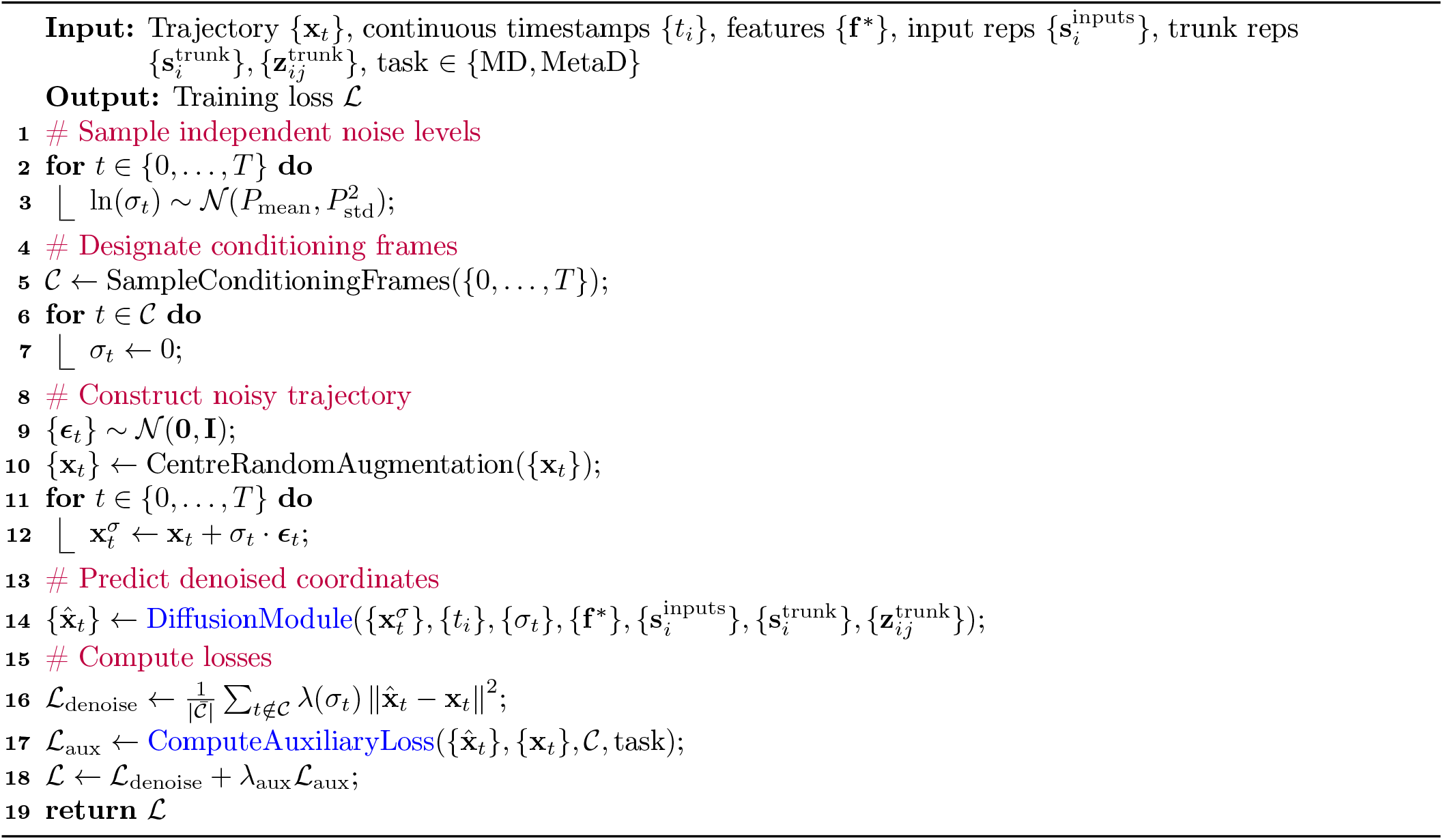

#### Algorithm 5 SampleDiffusion (Forecasting / Interpolation)

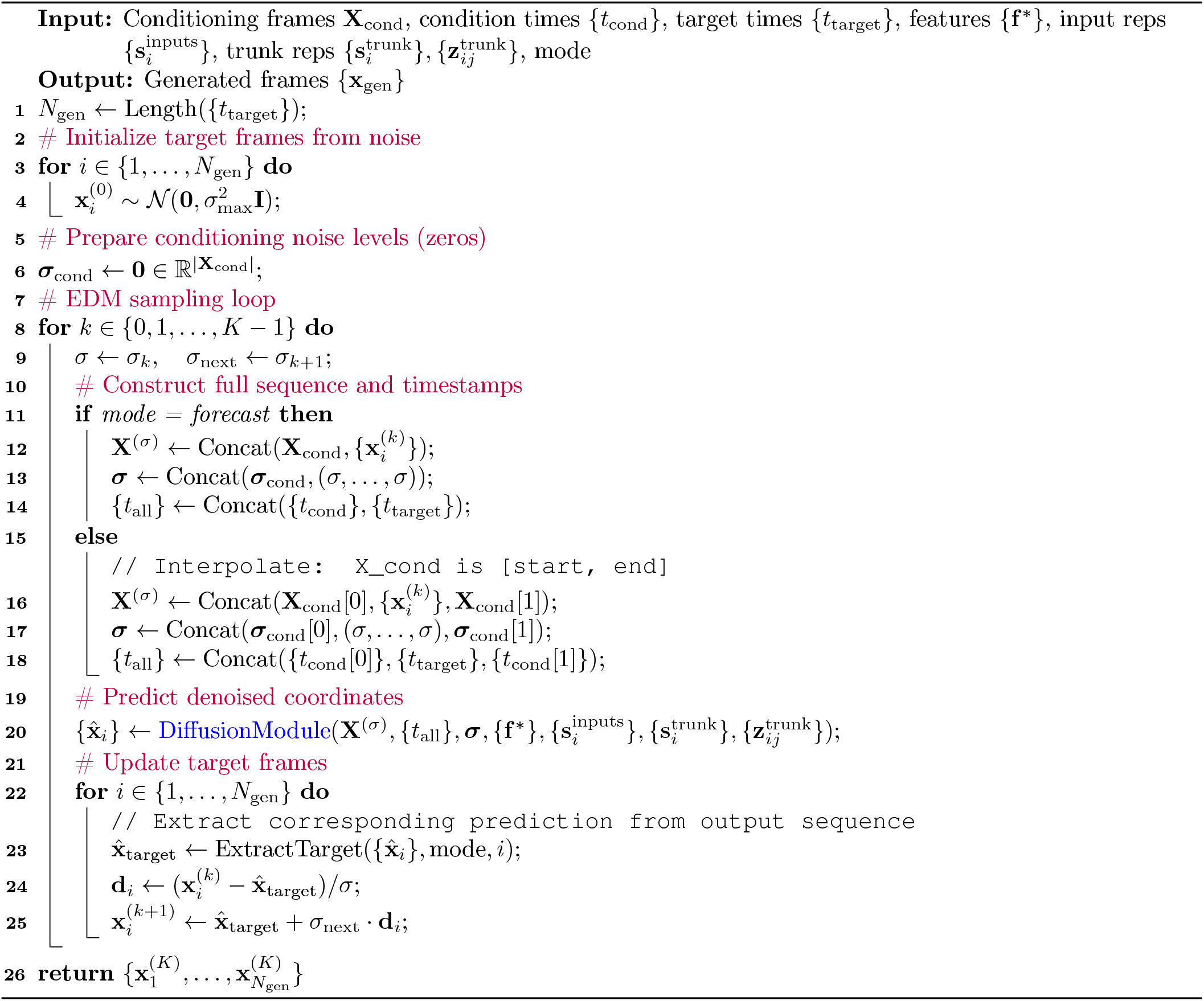

#### Algorithm 6 >Diffusion Module

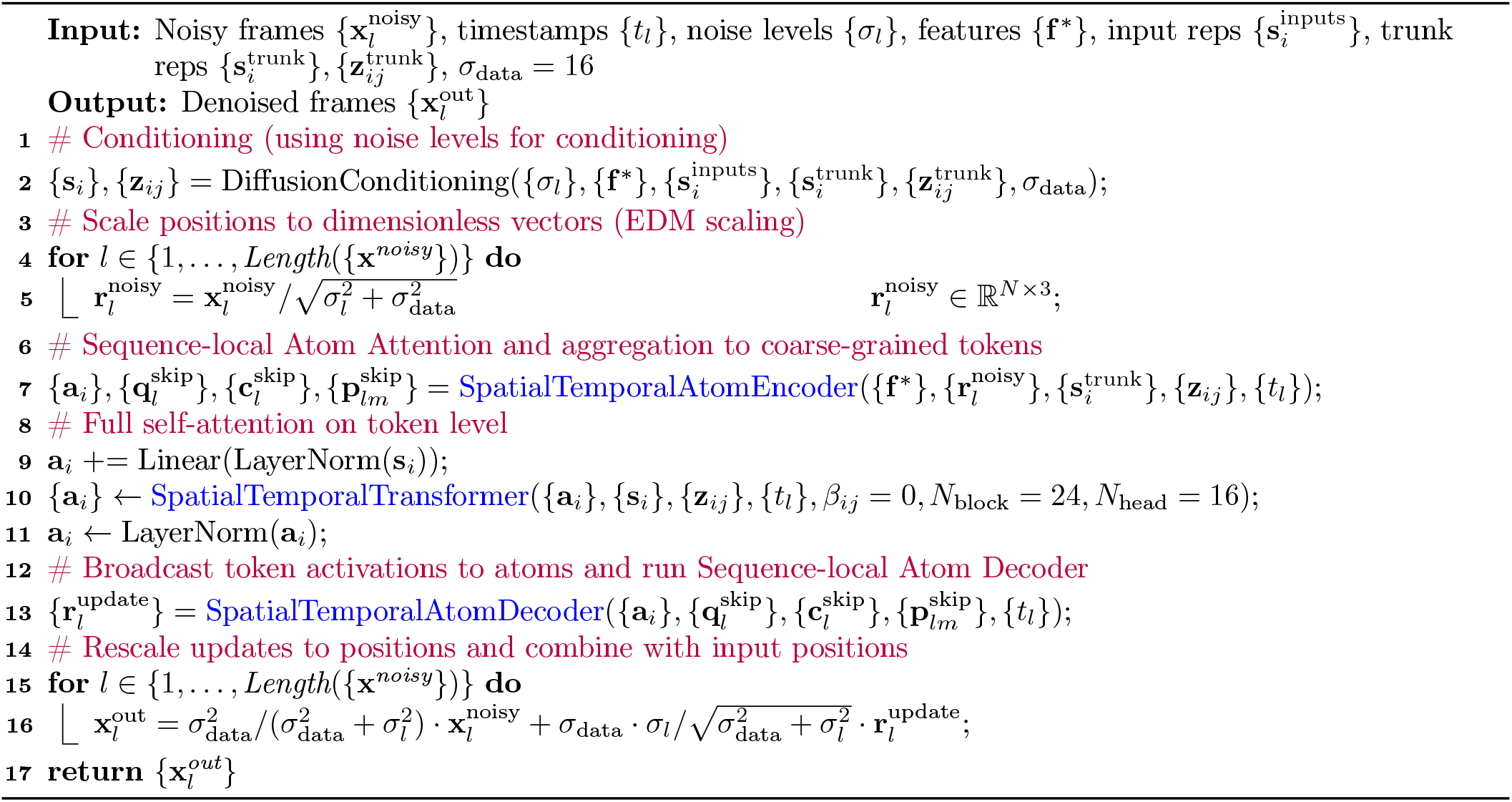

#### Algorithm 7 SpatialTemporalTransformer

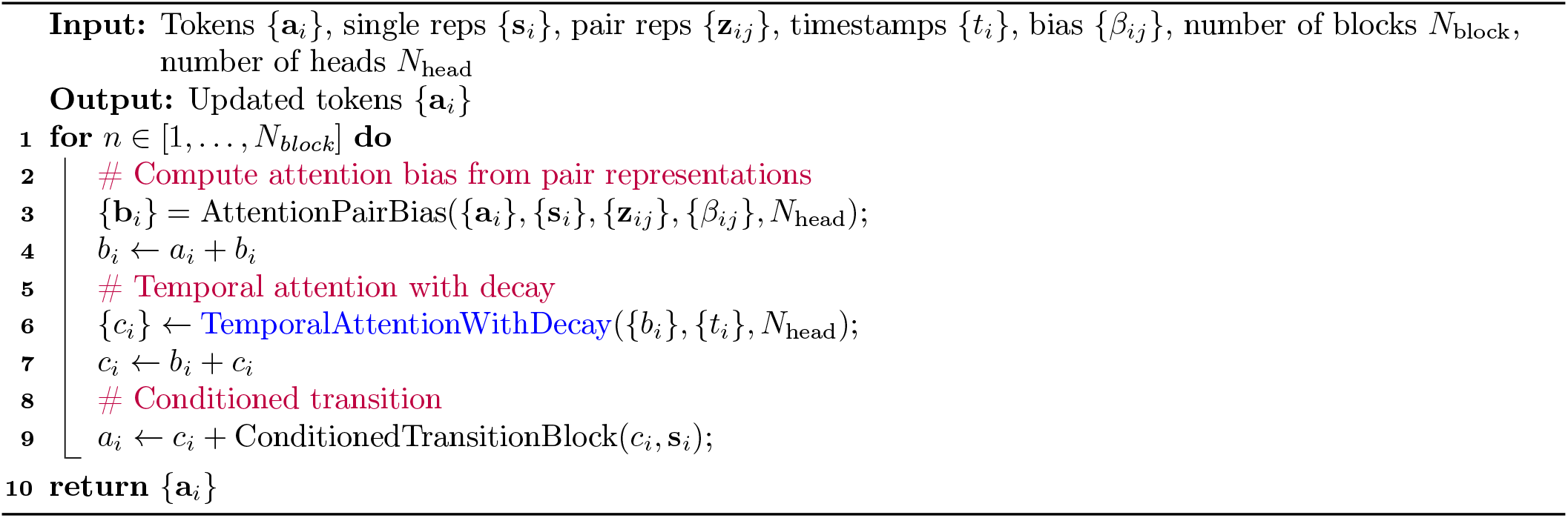

#### Algorithm 8 TemporalAttentionWithDecay

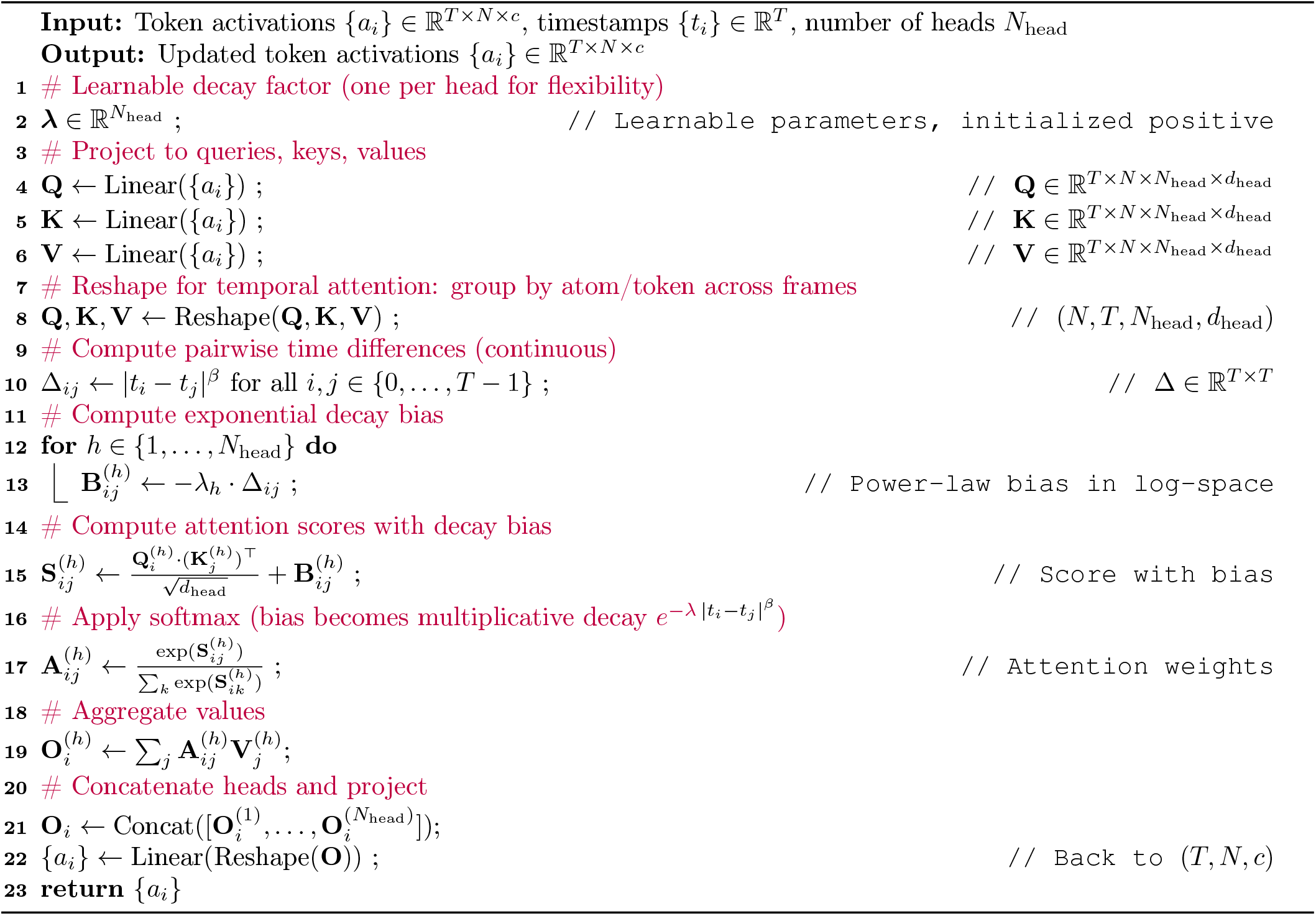

**Supplementary Figure 1.**
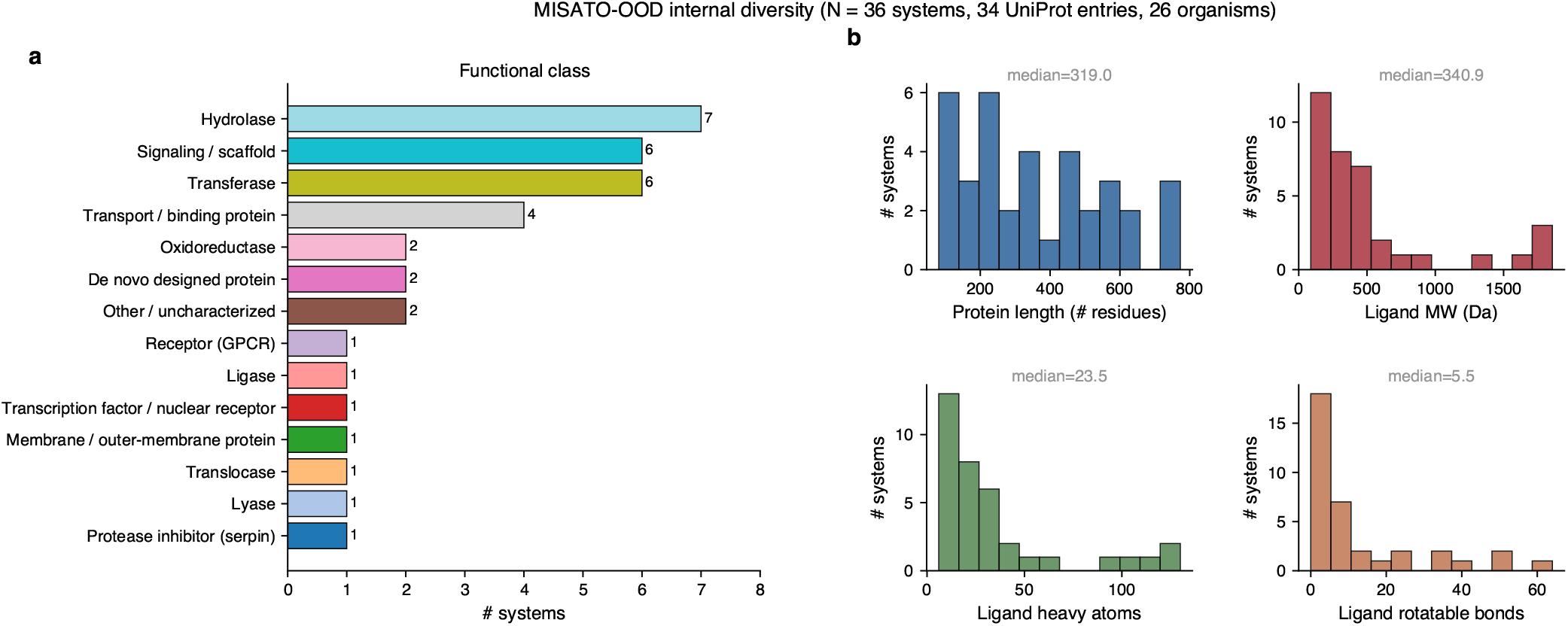
Internal diversity of the MISATO-OOD evaluation subset (N = 36 systems). (a) Coarse functional-class breakdown derived from EC numbers and UniProt annotation; the 36 systems span 34 distinct UniProt entries from 26 source organisms and cover all six top-level enzyme classes (16 enzymes) alongside non-enzymatic receptors, signaling/scaffold proteins, transporters and other proteins. (b) Distributions of protein length, ligand molecular weight, ligand heavy-atom count, and ligand rotatable-bond count.

**Supplementary Figure 2.**
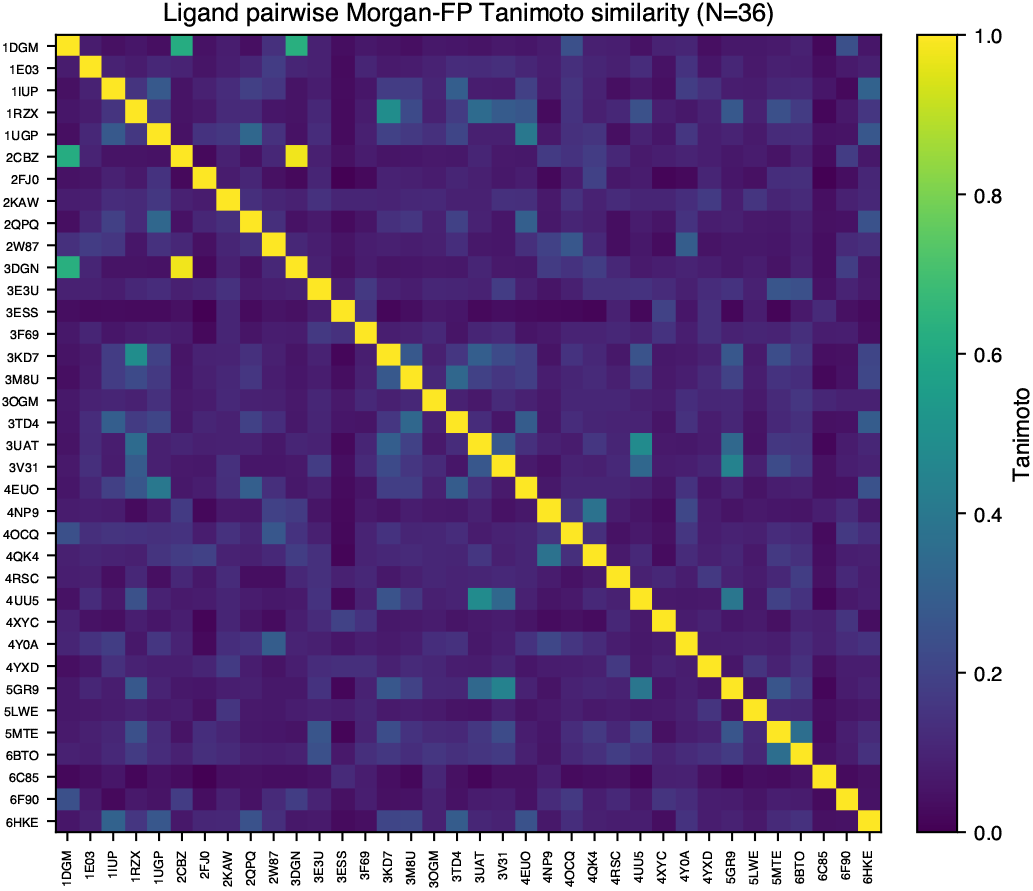
Internal diversity of the MISATO-OOD evaluation subset (N = 36 systems). Pairwise ligand Bemis–Murcko scaffold Morgan-fingerprint Tanimoto similarity (median 0.09), confirming chemically diverse ligands. Systems that fail at inference are excluded.

**Supplementary Figure 3.**
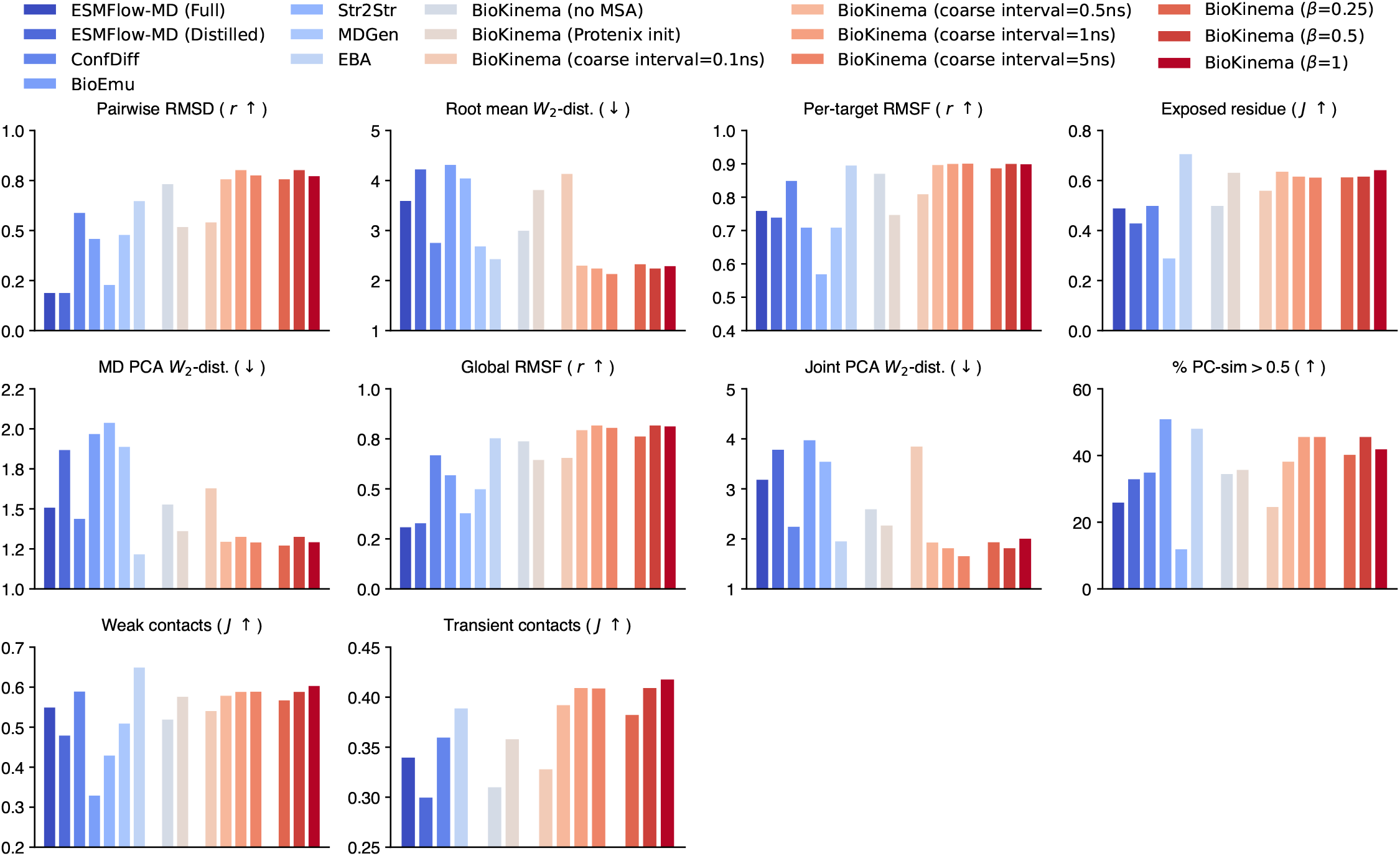
Bar plots comparing BioKinema and competing methods on the ATLAS test set across the 10 metrics from AlphaFlow^20^. Bars are grouped (left to right) into baselines, two BioKinema input ablations (*no MSA*, which removes the multiple-sequence-alignment input, and *Protenix init*, which initialises generation from a Protenix-predicted structure rather than an MD frame), BioKinema at coarse temporal intervals Δ*t* ∈ {0.1, 0.5, 1, 5} ns, and an ablation of the temporal-scaling exponent *β* ∈ {0.25, 0.5, 1}, which sets how the inter-frame time interval is encoded in the temporal attention. Values are aggregated across n = 81 targets. Unless otherwise stated, coarse interval is set to 1 ns and *β* = 0.5 by default.

**Supplementary Figure 4.**
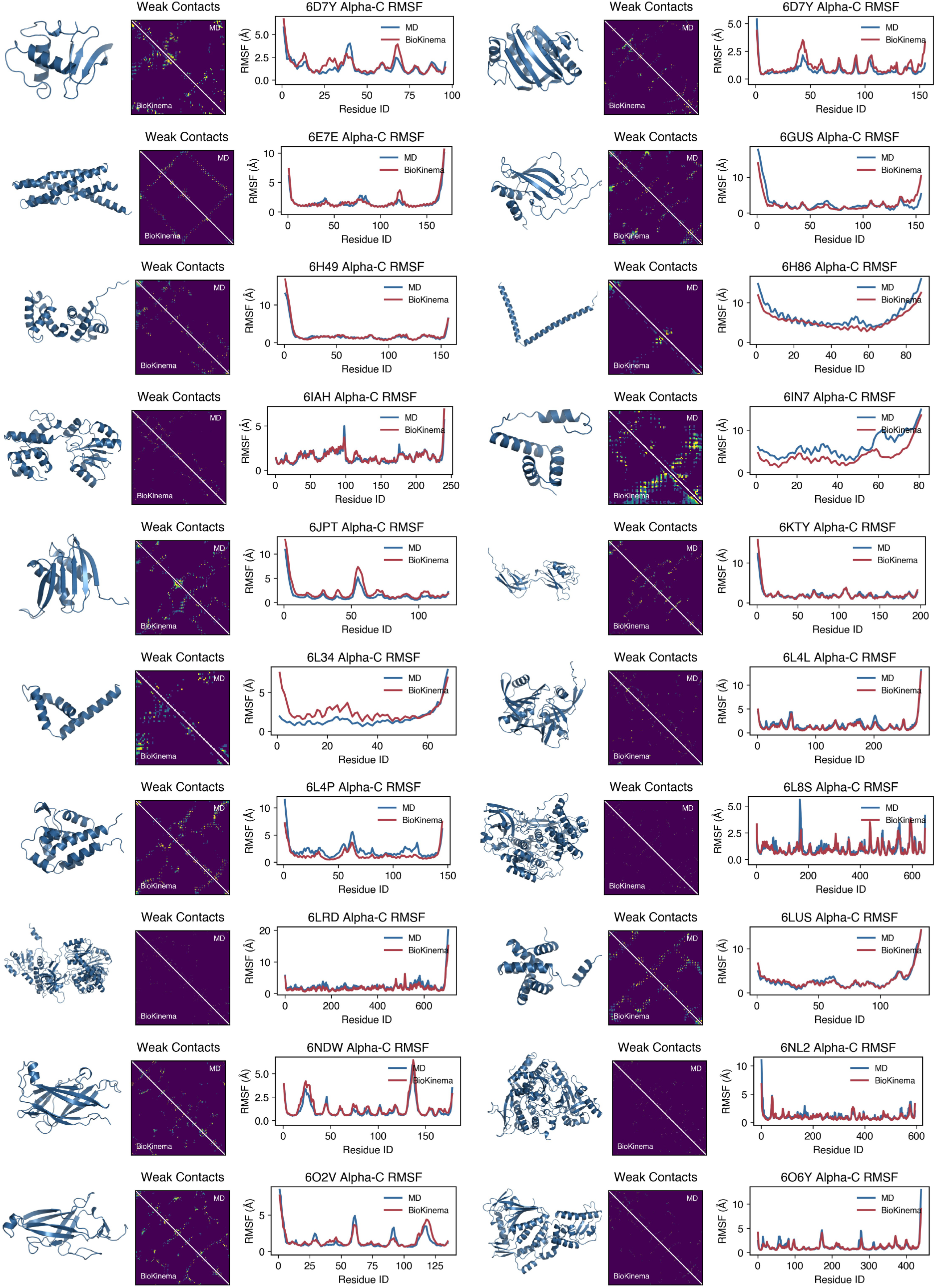
Results on the ATLAS-OOD test set (part 1/3). For each case, protein structures (left), weak contacts map (middle), and line plot showing Cα RMSF for both MD and BioKinema are shown. We define pairs of residues as contacted if the distance ≤ 8 Å. Weak contacts are defined as pairs of residues with contact probability ≤ 50%.

**Supplementary Figure 5.**
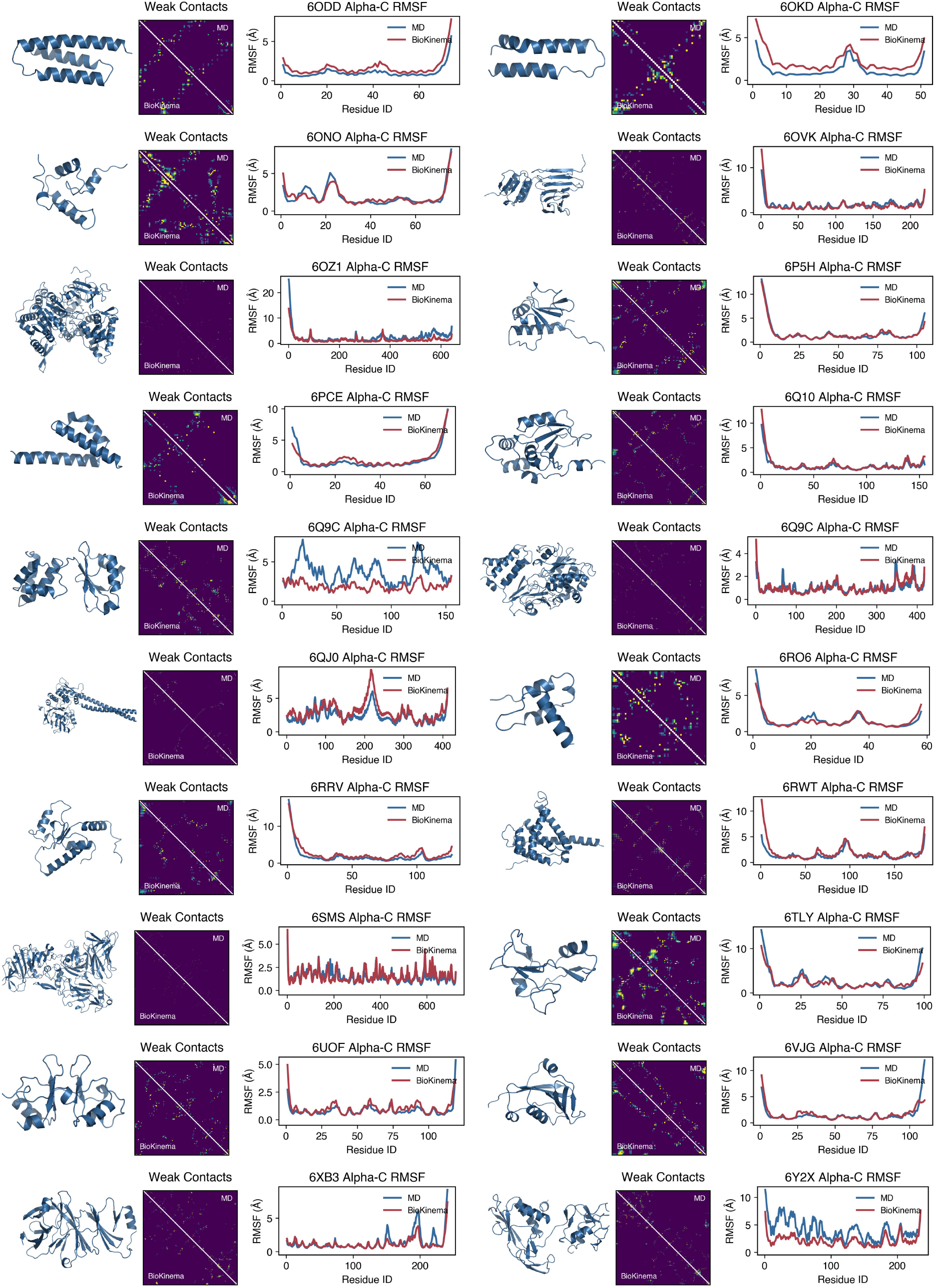
Results on the ATLAS-OOD test set (part 2/3). For each case, protein structures (left), weak contacts map (middle), and line plot showing Cα RMSF for both MD and BioKinema are shown. We define pairs of residues as contacted if the distance ≤ 8 Å. Weak contacts are defined as pairs of residues with contact probability ≤ 50%.

**Supplementary Figure 6.**
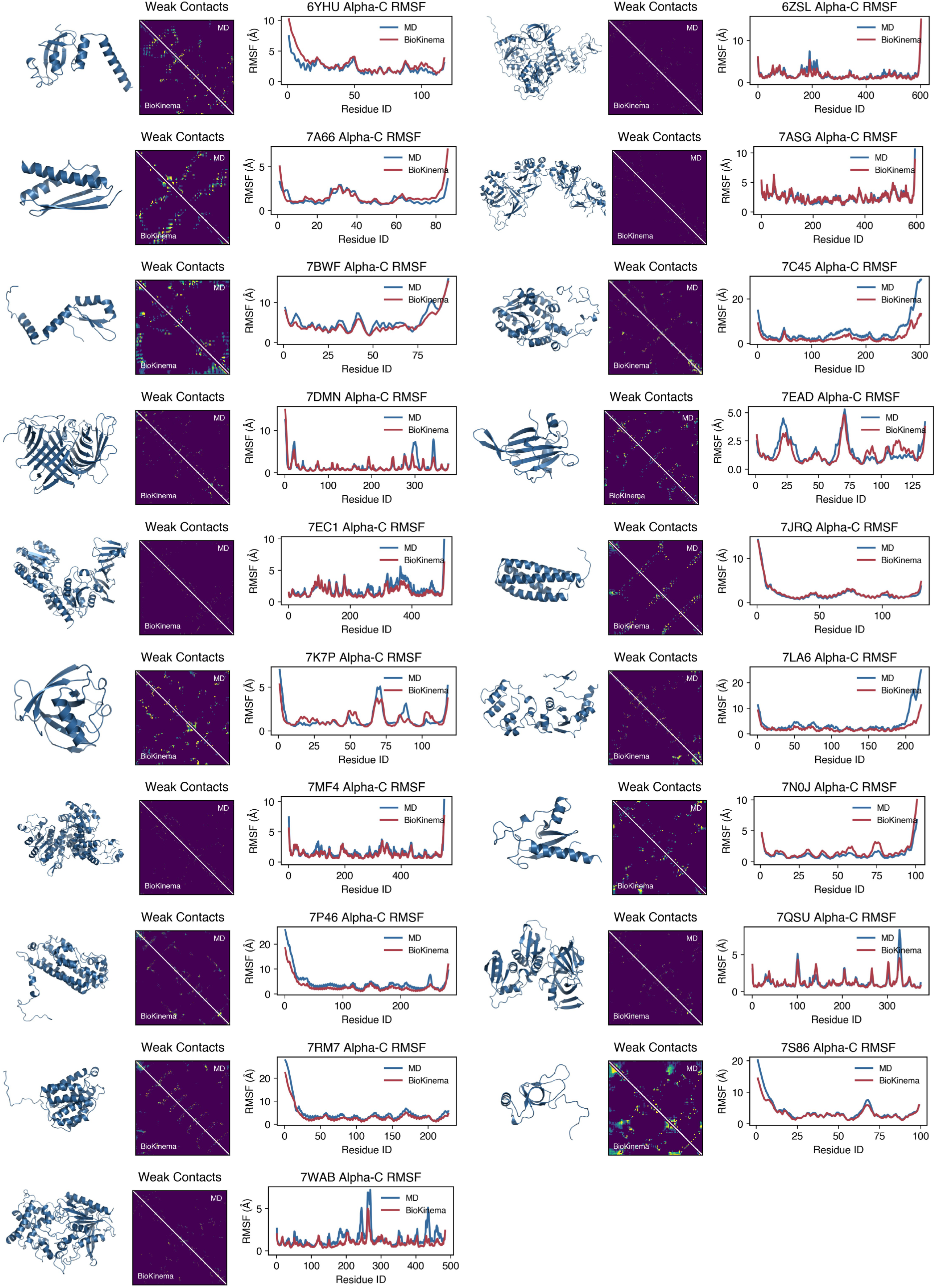
Results on the ATLAS-OOD test set (part 3/3). For each case, protein structures (left), weak contacts map (middle), and line plot showing Cα RMSF for both MD and BioKinema are shown. We define pairs of residues as contacted if the distance ≤ 8 Å. Weak contacts are defined as pairs of residues with contact probability ≤ 50%.

**Supplementary Figure 7.**
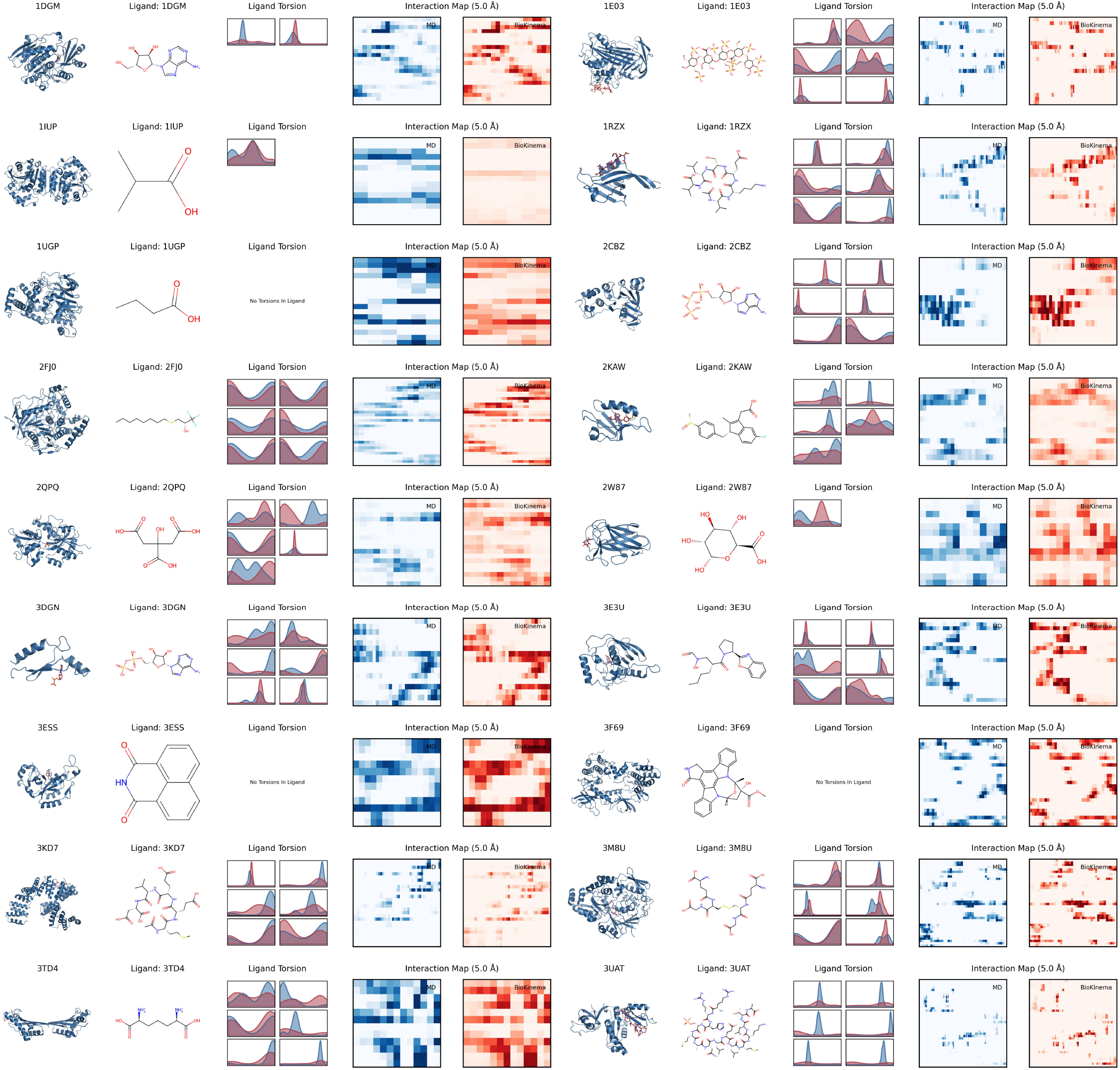
Results on the MISATO-OOD test set (part 1/2). For each case, complex structures (left), ligand torsions of up to 6 rotatable bonds (middle), and interaction maps for both MD and BioKinema are shown. We define “rotatable bond” as a non-ring single bond between heavy atoms. A residue is considered in contact with a ligand atom if any heavy atom of the residue is within 5.0 Å. The horizontal axis represents ligand heavy atoms (with names from the source SDF), and the vertical axis represents pocket residues.

**Supplementary Figure 8.**
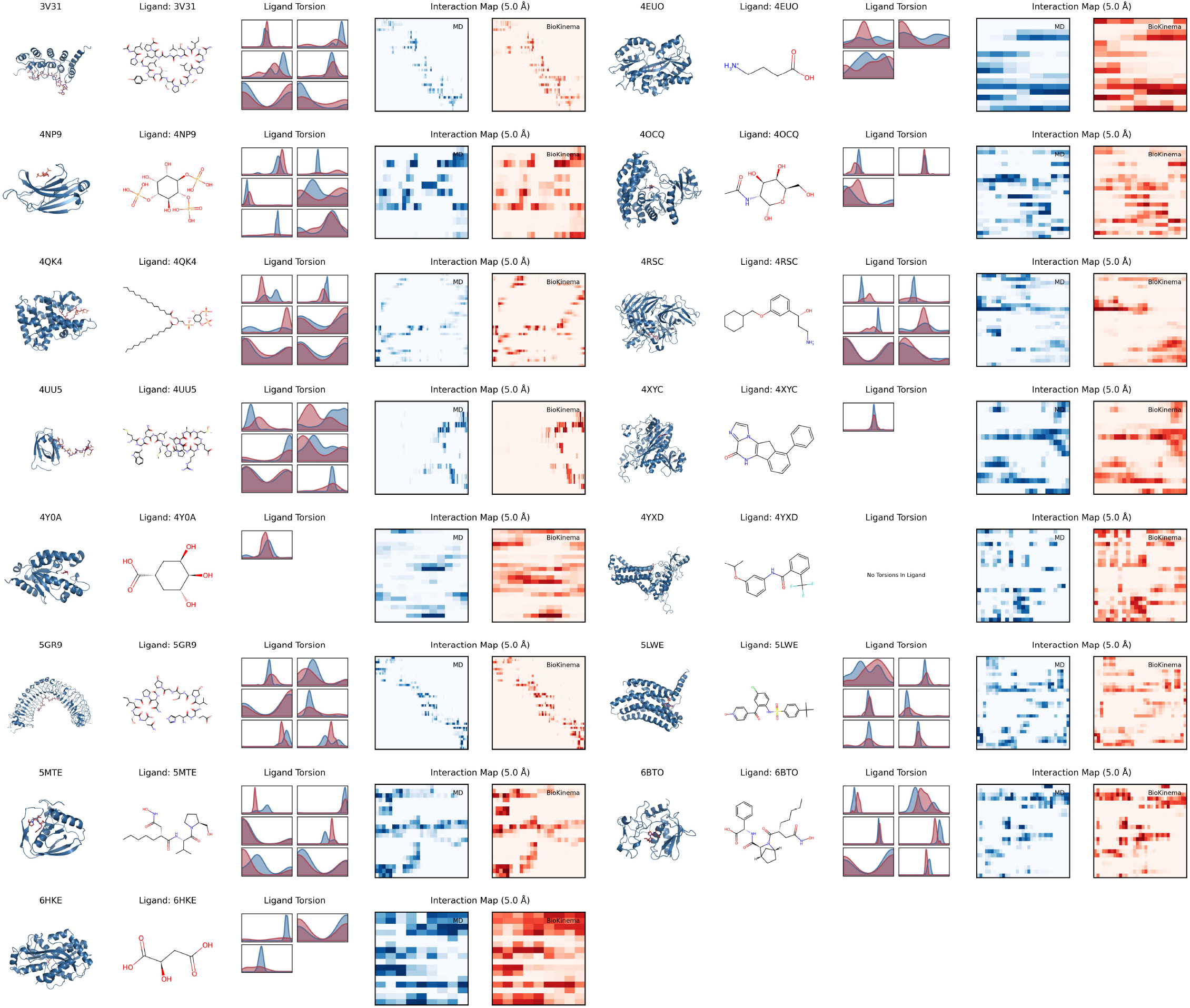
Results on the MISATO-OOD test set (part 2/2). For each case, complex structures (left), ligand torsions of up to 6 rotatable bonds (middle), and interaction maps for both MD and BioKinema are shown. We define “rotatable bond” as a non-ring single bond between heavy atoms. A residue is considered in contact with a ligand atom if any heavy atom of the residue is within 5.0 Å. The horizontal axis represents ligand heavy atoms (with names from the source SDF), and the vertical axis represents pocket residues.

**Supplementary Figure 9.**
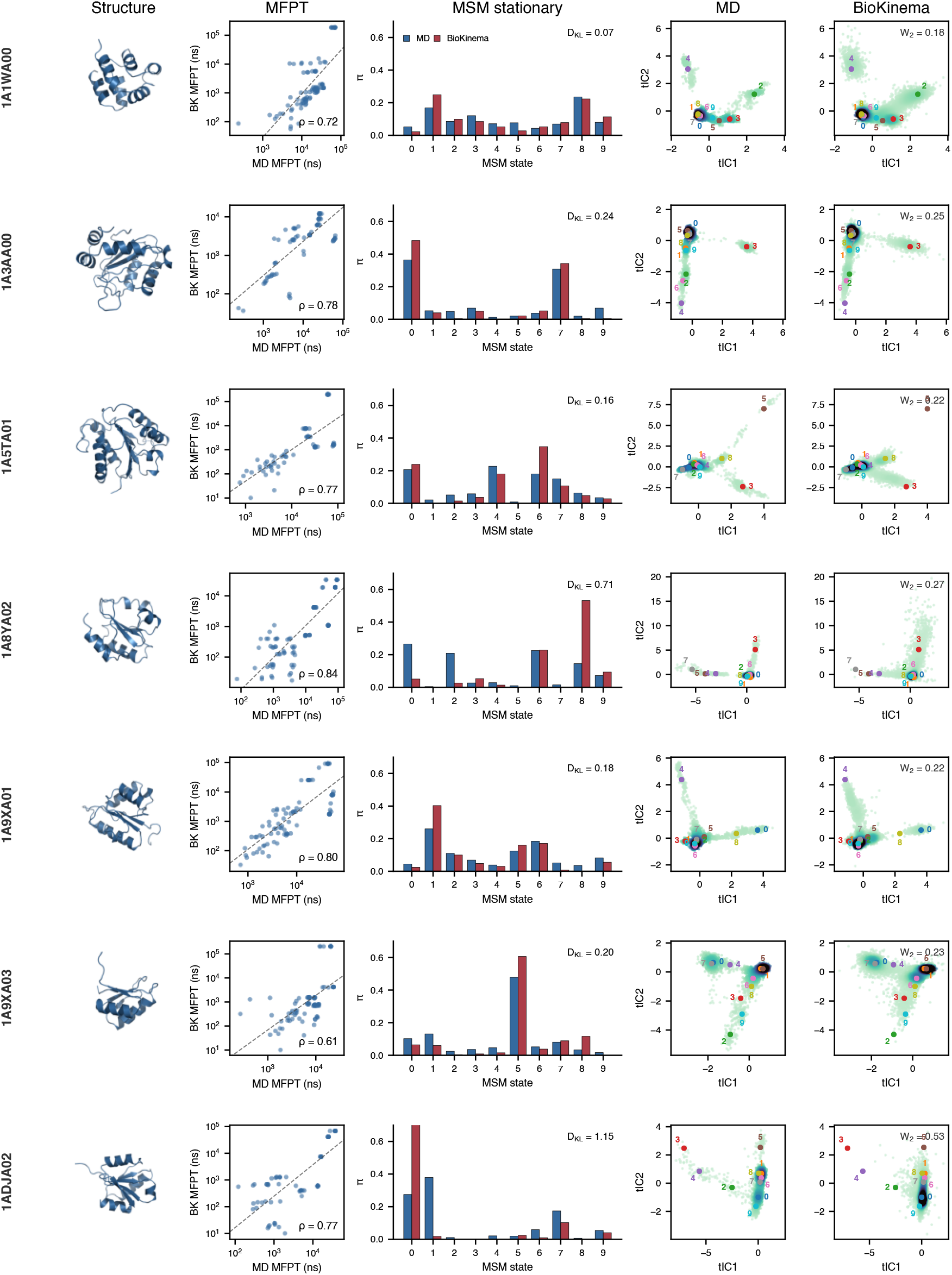
Per-system kinetics and thermodynamics on the CATH2 test set (1/3). Each column shows, from left to right, the protein structure, the per-protein MFPT scatter with log-space Pearson correlation *ρ* annotated, the MSM stationary distribution over *K*=10 states for MD (blue) and BioKinema (red) with *D*_KL_ annotated, and the pooled TICA two-dimensional density coloured by point density with the MSM cluster centres (colored circles, cluster id labelled) and *W*_2_ distance annotated.

**Supplementary Figure 10.**
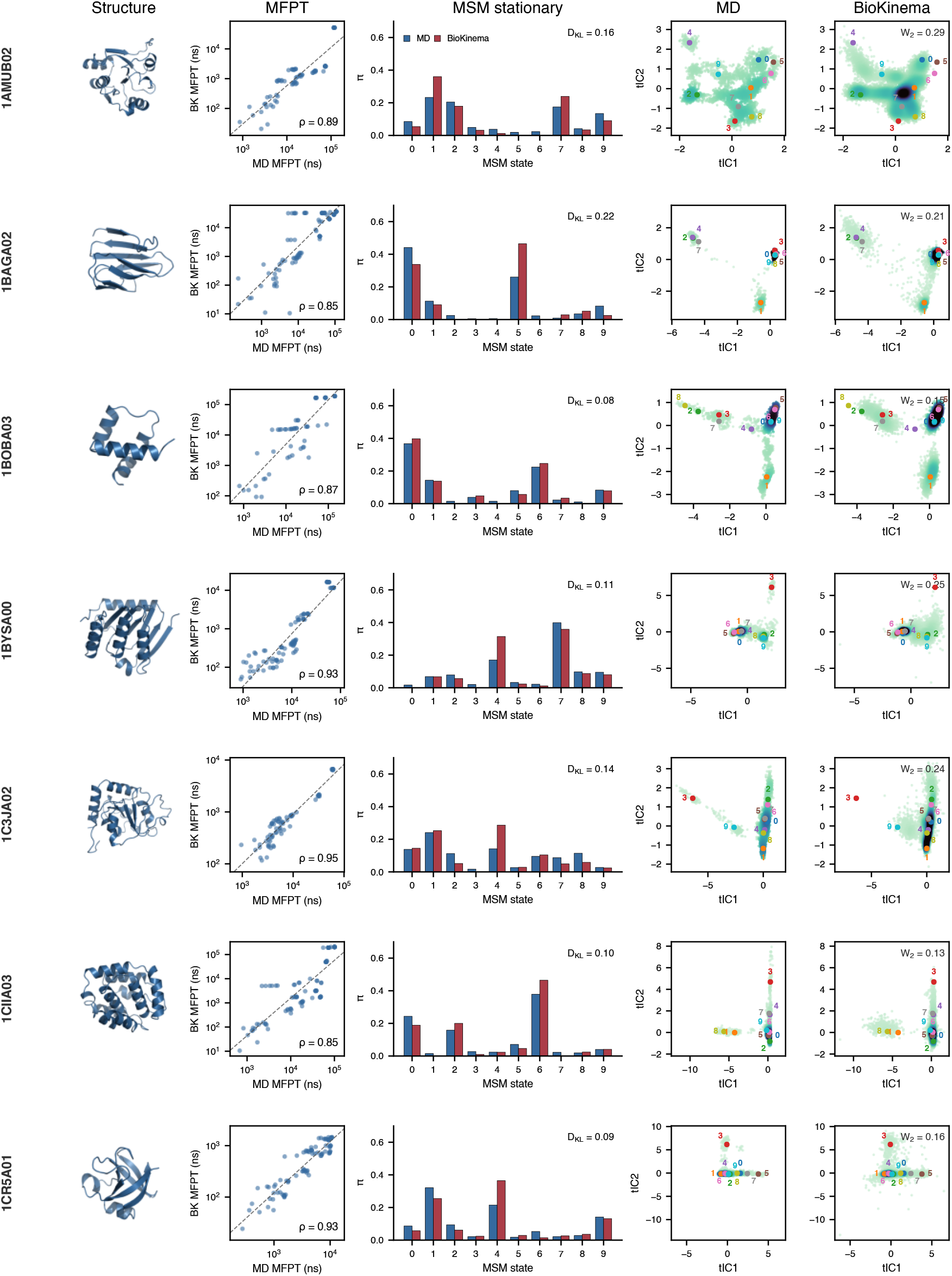
Per-system kinetics and thermodynamics on the CATH2 test set (2/3). Each column shows, from left to right, the protein structure, the per-protein MFPT scatter with log-space Pearson correlation *ρ* annotated, the MSM stationary distribution over *K*=10 states for MD (blue) and BioKinema (red) with *D*_KL_ annotated, and the pooled TICA two-dimensional density coloured by point density with the MSM cluster centres (colored circles, cluster id labelled) and *W*_2_ distance annotated.

**Supplementary Figure 11.**
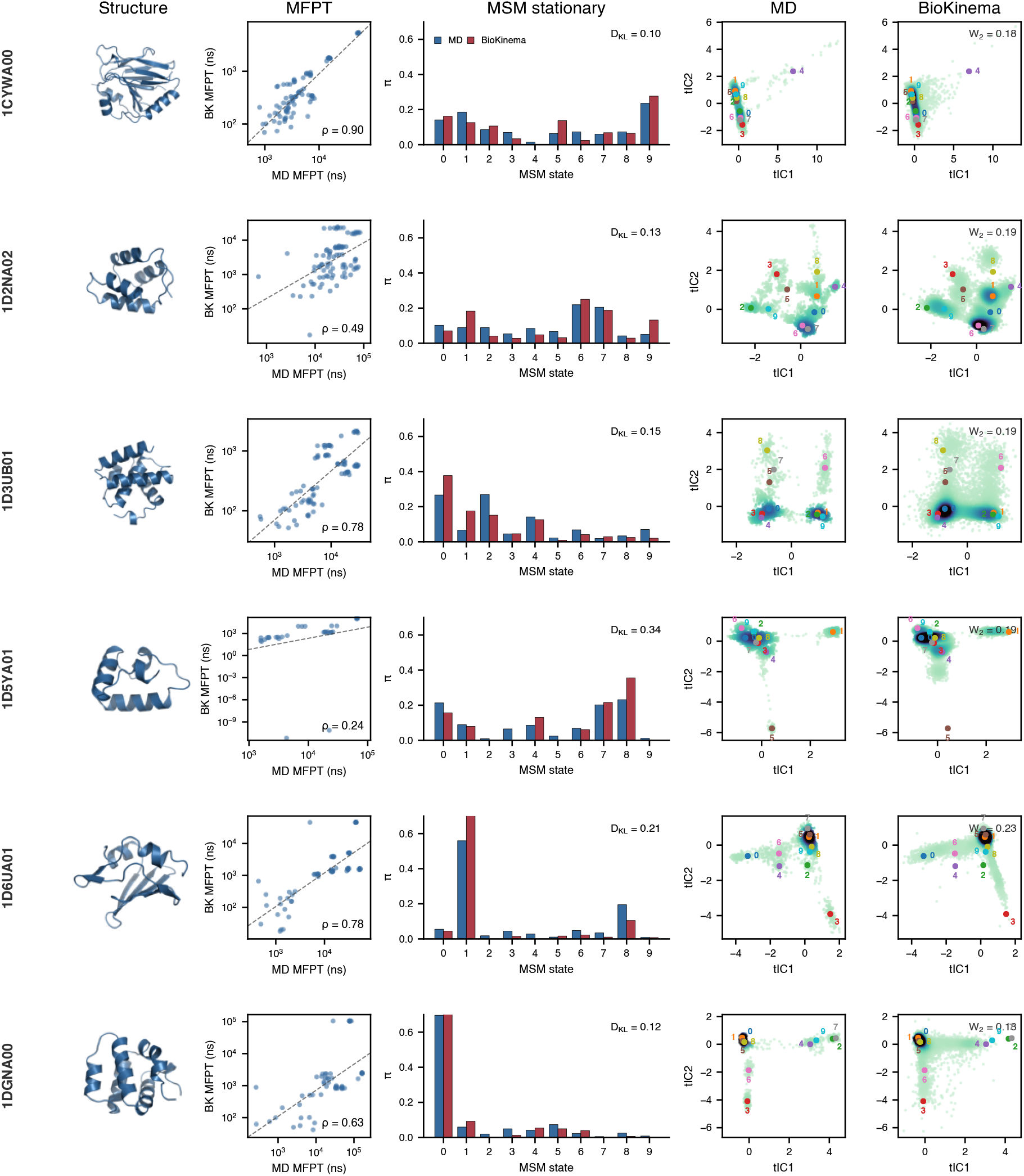
Per-system kinetics and thermodynamics on the CATH2 test set (3/3). Each column shows, from left to right, the protein structure, the per-protein MFPT scatter with log-space Pearson correlation *ρ* annotated, the MSM stationary distribution over *K*=10 states for MD (blue) and BioKinema (red) with *D*_KL_ annotated, and the pooled TICA two-dimensional density coloured by point density with the MSM cluster centres (colored circles, cluster id labelled) and *W*_2_ distance annotated.

**Supplementary Figure 12.**
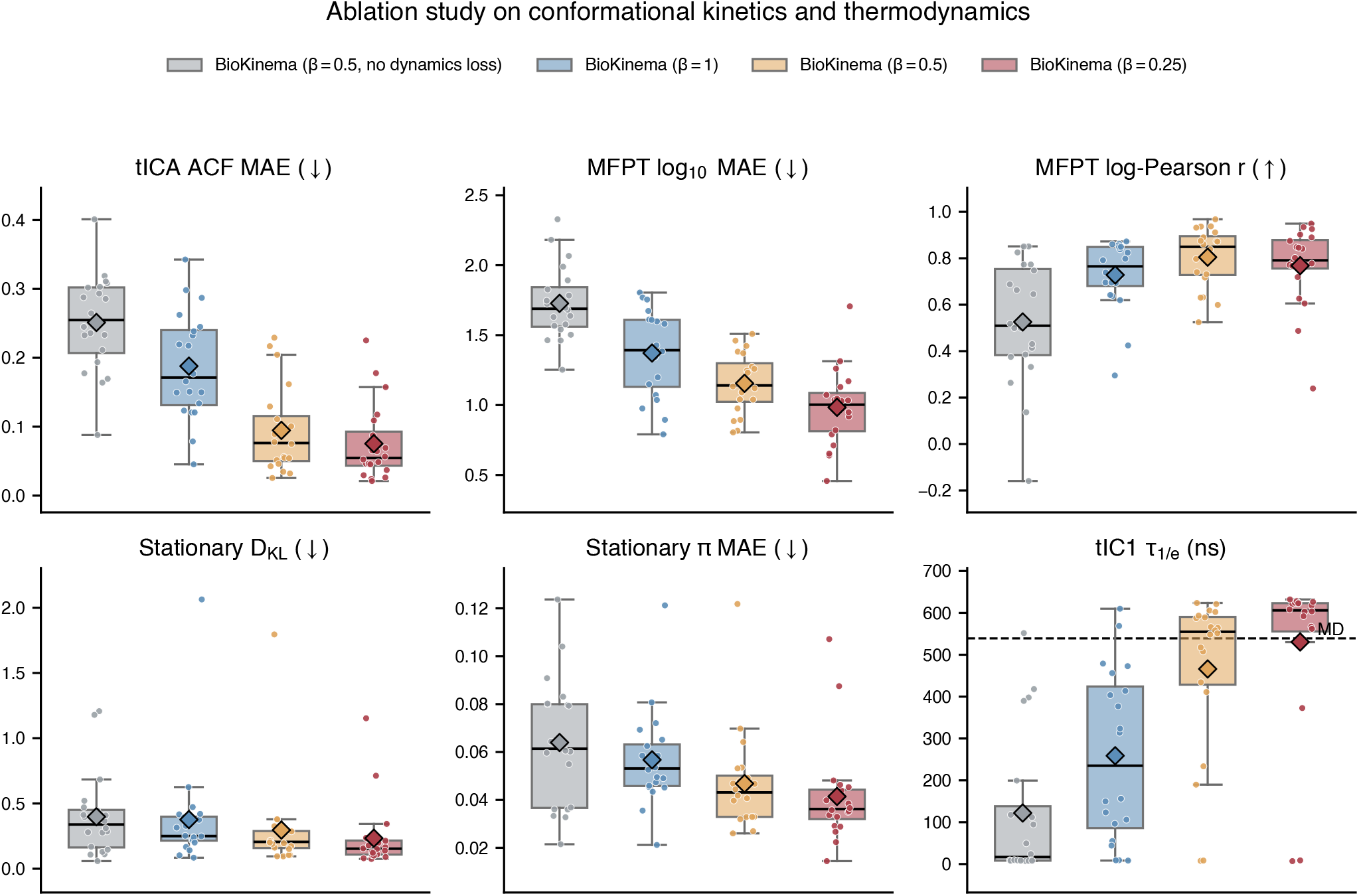
Ablation of the time-scaling exponent and the TICA–MSM dynamics loss on conformational kinetics and thermodynamics. Box plots over the 20 CATH2-OOD test systems comparing three models trained with different *β* ∈ {1, 0.5, 0.25}, and one model trained with *β*=0.5 but without the dynamics loss. Five 1 *µ*s trajectories at 10 ns resolution are generated from each MD initial frame. Panels show, for kinetics, the TICA autocorrelation-function MAE, the log_10_(MFPT) MAE, and the log-space MFPT Pearson correlation; and, for thermodynamics, the stationary-distribution *D*_KL_, the per-state stationary MAE, and the tIC1 relaxation time *τ*_1/e_ (dashed line, MD reference).

**Supplementary Figure 13.**
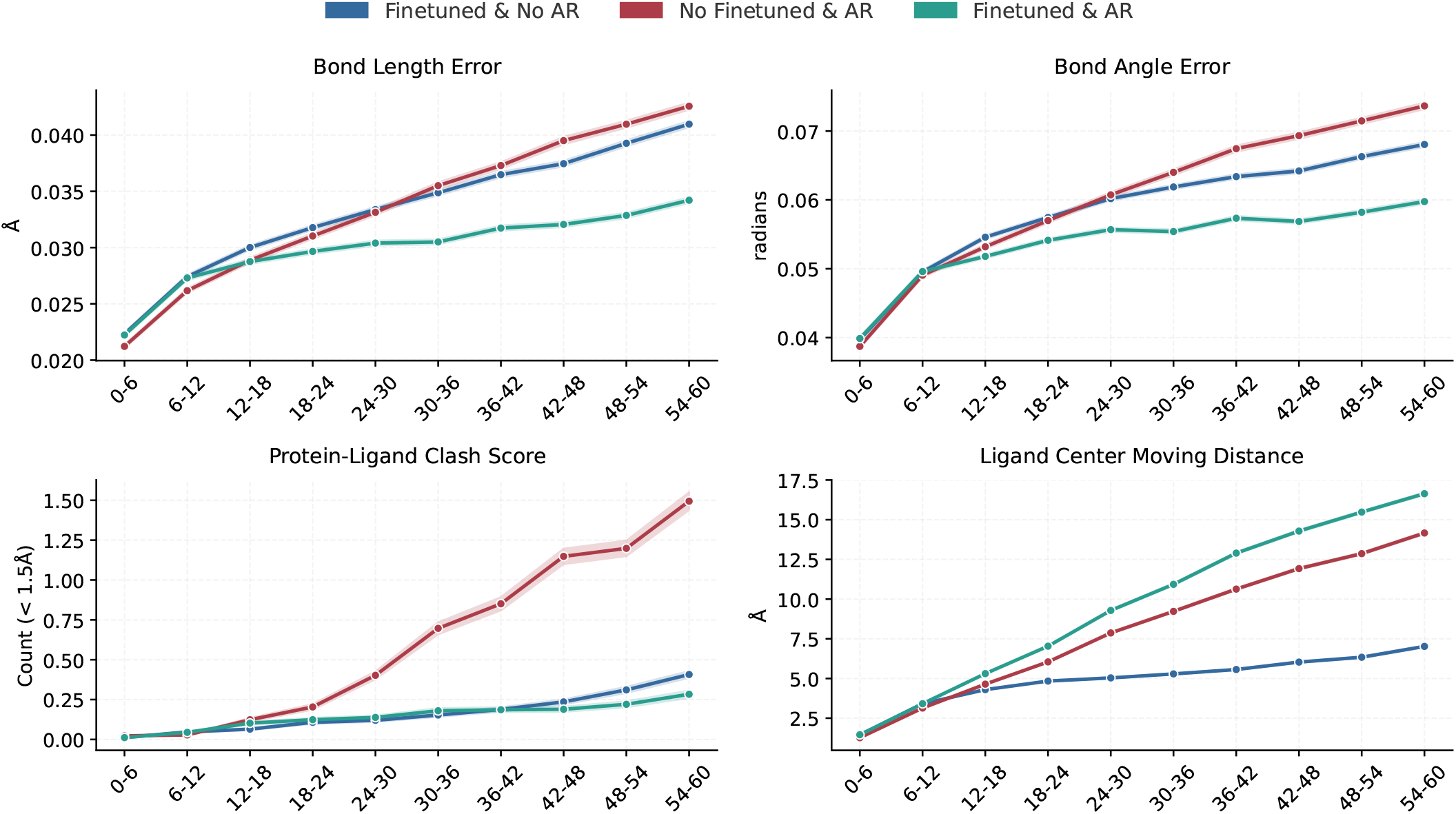
Line plot comparing different versions of our model in terms of bond length error, bond angle error, protein-ligand clash score, and ligand center moving distance across 60 ps trajectories, evaluated on 31 protein-ligand systems from the DD-13M test set. “Finetuned” denotes that the model is finetuned from BioKinema and “AR” denotes the auto-regressive method. All trajectories are generated 20 times independently. The auto-regressive method adopts a 10 ps time step for each iteration.

**Supplementary Figure 14.**
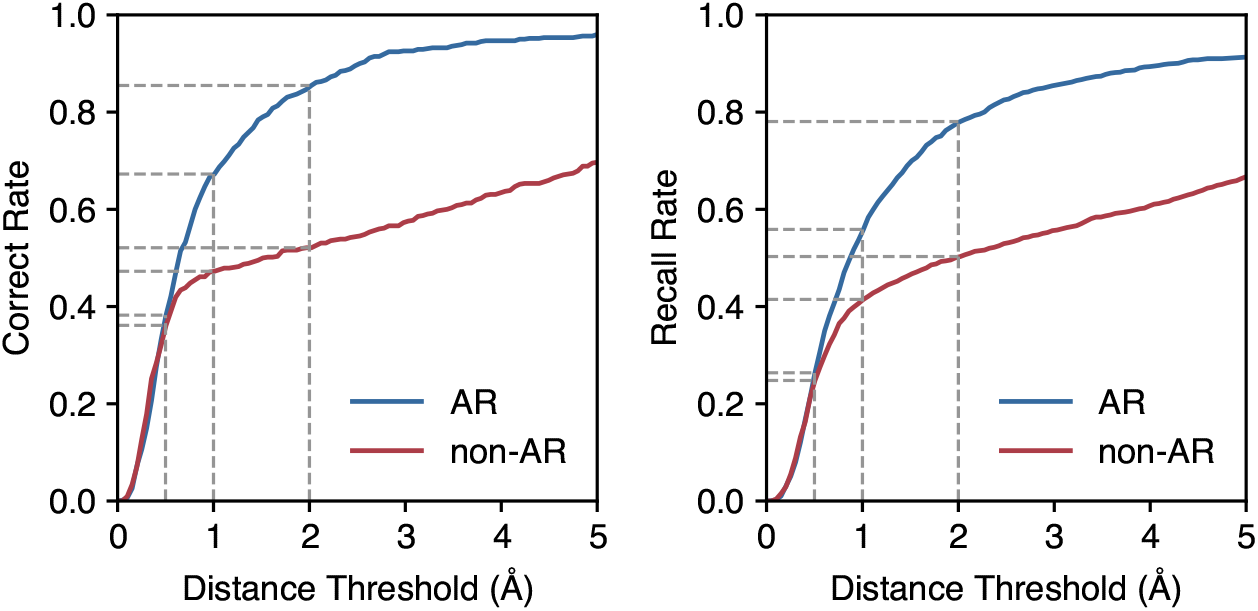
Line plot showing precision (correct rate) and recall curves as a function of endpoint distance threshold comparing auto-regressive (blue) and non-auto-regressive (red) methods, evaluated on 31 protein-ligand systems from the DD-13M test set. All models are finetuned from BioKinema. All trajectories are generated 20 times independently. The auto-regressive method adopts a 10 ps time step for each iteration.

**Supplementary Figure 15.**
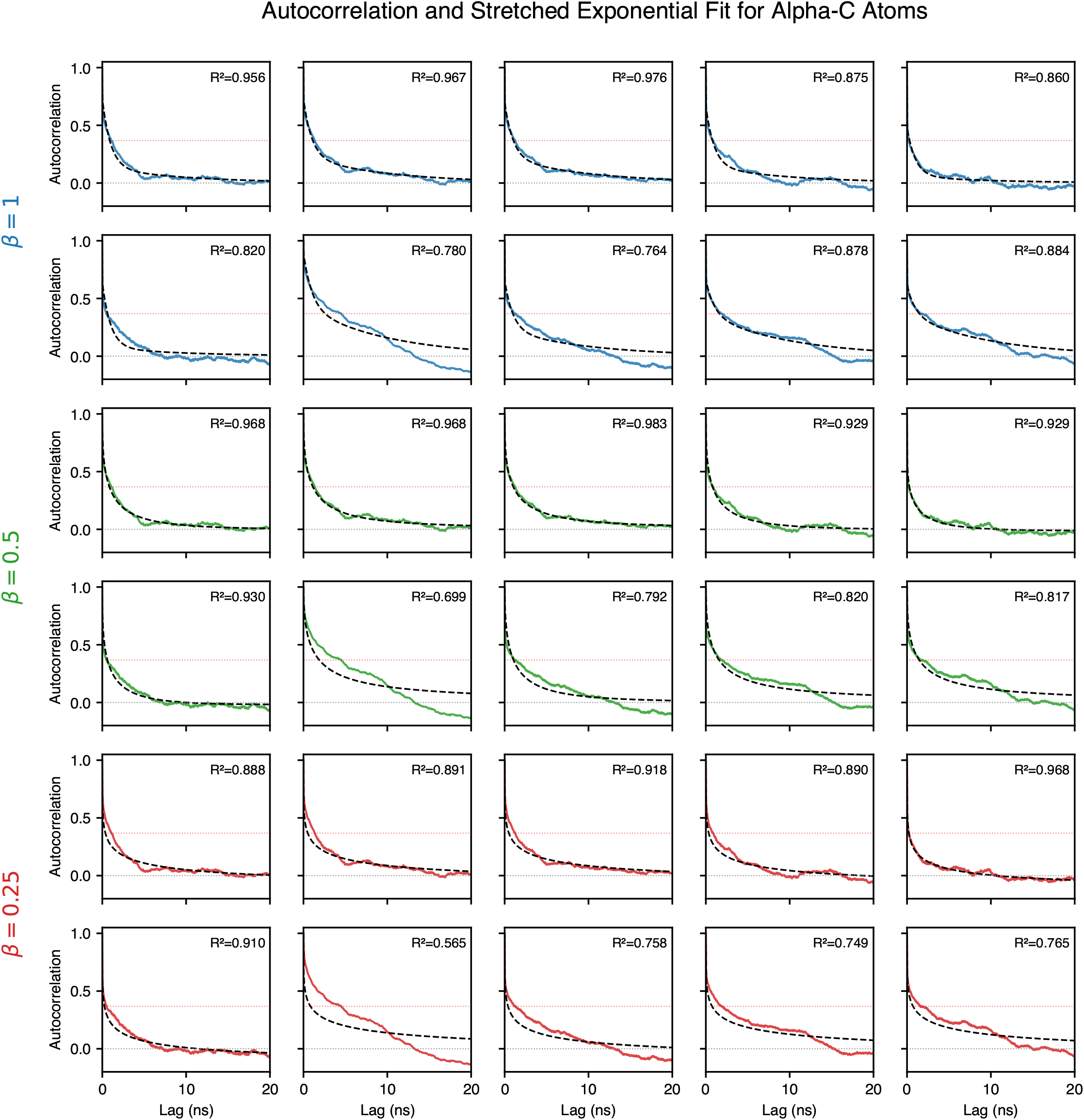
Line plots showing autocorrelation curves of C*α* atom coordinates and their fitted combined stretched-exponential curves. The combined exponential is defined as the weighted combination of four exponential decays 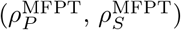 with different decay factors *λ* (0.01, 0.1, 1, and 10).

